# The N-terminal helix of MarA as a key element in the mechanism of DNA binding

**DOI:** 10.1101/2024.02.13.580091

**Authors:** Marina Corbella, Cátia Moreira, Roberto Bello-Madruga, Marc Torrent, Shina C.L. Kamerlin, Jessica M.A. Blair, Enea Sancho-Vaello

## Abstract

Efflux is one of the mechanisms employed by Gram-negative bacteria to become resistant to routinely used antibiotics. The inhibition of efflux by targeting their regulators is a promising strategy to re-sensitise bacterial pathogens to antibiotics. AcrAB-TolC is the main Resistance-Nodulation-Division efflux pump in Enterobacteriaceae. MarA is an AraC/XylS family global regulator that regulates more than 40 genes related to the antimicrobial resistance phenotype, including *acrAB*. The aim of this work was to understand the role of the N-terminal helix of MarA in the mechanism of DNA binding. An N-terminal deletion of MarA showed that the N-terminal helix has a role in the recognition of the functional marboxes. By engineering two double cysteine variants of MarA, and combining *in vitro* electrophoretic mobility assays and *in vivo* measurements of *acrAB* transcription with molecular dynamic simulations, it was shown that the immobilization of the N-terminal helix of MarA prevents binding to DNA. This new mechanism of inhibition seems to be universal for the monomeric members of the AraC/XylS family, as suggested by additional molecular dynamics simulations of the two-domain protein Rob. These results point to the N-terminal helix of the AraC/XylS family monomeric regulators as a promising target for the development of inhibitors.

## Introduction

Antimicrobial resistance (AMR) is a silent pandemic with 1.27 million deaths directly attributable to resistance in 2019 ^1,2^. In light of this scenario, urgent action is required including strong action on infection prevention and control programmes, the search for new antibiotics and the search for “resistance breaker” strategies to re-sensitise bacteria to existing antibiotics. Basic research to increase the understanding of mechanisms underpinning antibiotic resistance are also critically important.

One of the mechanisms employed by Gram-negative bacteria to become resistant to routinely used antibiotics is the use of efflux pumps to export antimicrobials out of bacterial cells ^3^. These efflux pumps primarily allow the microorganisms to regulate their internal environment by removing toxic substances, but other physiological roles have also been proposed, including the transport of antimicrobial peptides or fatty acids, cell to cell signalling, virulence and bacterial biofilm formation ^4,5^. There are seven families of efflux pumps, with the members of the Resistance-Nodulation-Division (RND) superfamily being the most clinically associated with MDR phenotypes ^4,6,7^. The members of the RND efflux pump superfamily form a tripartite assembly consisting of a trimeric outer membrane protein channel belonging to the outer membrane factor (OMF) family, a hexameric periplasmic adaptor protein (PAP) and a trimeric inner membrane protein ^7–12^. Some RND efflux pumps include an accessory protein such as AcrZ in the case of the AcrAB–TolC efflux pump ^13^. Members of the RND superfamily include the *E. coli* AcrAB–TolC, MexAB-OprM in *Pseudomonas*, MtrCDE in *Neisseria gonorrhoeae*, and *Acinetobacter* Ade systems (AdeABC and AdeIJK) ^7,8,14,15^.

AcrAB-TolC is the main RND efflux pump in Enterobacteriaceae including *Escherichia coli, Salmonella enterica* and *Klebsiella pneumoniae.* The regulation of the expression of this efflux pump is complex and includes global regulators such as MarA, SoxS and Rob, which increases the expression of the major efflux genes ^16–18^. Additionally, MarA regulates 40 other genes, many of them related to the antimicrobial resistance phenotype ^16–18^. MarA, SoxS and Rob share a high percentage of identity and similarity (e.g., SoxS and MarA share 41% similarity and 67% identity ^19^), and the N-terminal DNA binding domain of Rob has 51% sequence identity and 71% sequence similarity with MarA ^20^). MarA, SoxS and Rob bind the same 20-bp highly degenerate DNA sequence, called the marbox. This degenerate sequence can be found in more than 10,000 locations in the *E. coli* genome, although most of them are non-functional ^21,22^.

MarA is encoded by the second gene in the *marRAB* operon ^18^. This operon is repressed by the dimeric MarR which binds to two specific palindromic sequences located in the promoter region *marO* ^23^. When a ligand, such as sodium salicylate, binds MarR, it releases the DNA and *marA* is transcribed activating or repressing the transcription of the genes under its control ^24^. MarA also controls the *marRAB* expression by binding as a monomer to the marbox located in the promoter region of the *marRAB* operon ^25^. When the signal disappears, MarR binds the DNA again and the repression resumes. Another layer of regulation is related to the inherent instability of MarA, which is associated with its degradation by the Lon protease ^26^. The *marA* gene encodes MarA, a monomeric protein exclusively composed of α-helices (Figure 1A). Helices 1-3 and 5-7 form the N- and C-terminal helix-turn-helix (HTH) domains, respectively, with helix 3 and 6 being in direct contact with the DNA ^27–29^. In the crystal structure, MarA bends the DNA by 35° to permit both HTH motifs to insert into the major groove simultaneously ^27^.

**Figure 1:**
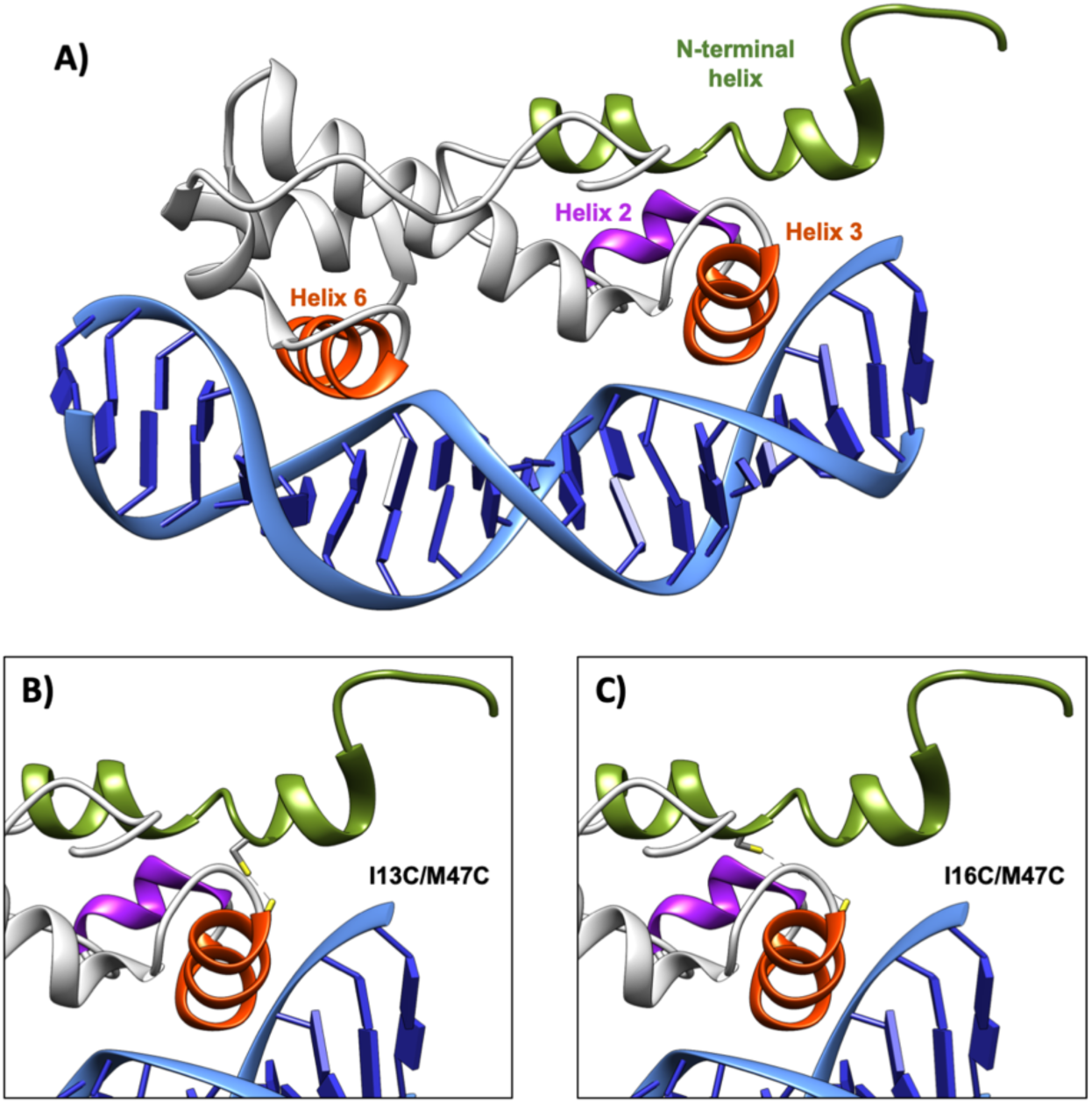
Structure of the WT MarA and its double cysteine variants used in this work. **(A)** MarA (PDB 1XS9) ^29,36^ is a monomer with two helix-turn-helix motifs. Helices 3 and 6 (in orange) interact directly with the major groove of the DNA. The N-terminal helix is shown in green. **(B)** and **(C)** Predicted structural models for the I13C/M47C and I16C/M47C variants. The formation of the disulfide bond in (B) the I13C/M47C variant maintains the N-terminal helix in the same position as observed in the PDBs, (C) while in I16C/M47C pulls it to a different position.

Several studies have suggested an important role for the N-terminal helix of MarA. By NMR, it was shown that the start of the helix is close to the DNA, suggesting that the N-terminal tail could contact the DNA backbone or help in the correct positioning of the helix 3 in the major groove ^28^. By mutagenesis, it was shown that the N-terminal helix was important for DNA binding ^30,31^. Also, the mutagenesis of 4 consecutive residues to alanine (L10-W13 and I14 to H17) increased the half-life and decreased the percentage of activation of both, the *zwf* and *fumC* promoters ^32^. By molecular dynamic simulations, it was shown that the flexibility of half of the N-terminal HTH motif dominates the motions of both MarA and Rob, showing the importance of the first nucleotides of the marbox (called A-box) for tight binding ^20,33^. In addition, the coexistence of meta-stable forms of MarA in solution or forming complexes with DNA has been shown ^28,34^. As it was shown that MarA exhibits multiple energetically similar conformations, it was postulated that residues distant from the active site can alter binding affinity and ligand specificity ^34^.

Many compounds that inhibit the action of efflux pumps (such as Phenylalanine-Arginine β-Naphthylamide (PAβN)) have been discovered, but the use of many is limited by their toxicity to human cells ^35^. An alternative strategy would be to target the regulators of the efflux pumps to prevent transcription of the pump genes. To date, there are no inhibitors for monomeric regulators belonging to the AraC/XylS family, such as MarA. The aim of this work is to understand the role of the N-terminal helix of MarA in the mechanism of DNA binding. An N-terminal deletion of MarA showed that the N-terminal helix has a role in the recognition of the functional marboxes. By engineering two double cysteine variants of MarA, and combining *in vitro* electrophoretic mobility assays and *in vivo* measurements of *acrAB* transcription with molecular dynamic simulations, it was shown that the immobilization of the N-terminal helix of MarA prevents binding to DNA. This new mechanism of inhibition seems to be universal for the monomeric members of the AraC/XylS family, as suggested by complementary molecular dynamics simulations of the two-domain protein Rob. These results point to the N-terminal helix of the AraC/XylS family monomeric regulators as a promising target for the development of inhibitors.

## Results

### The N-terminal helix of MarA is a crucial structural element for the mechanism of DNA binding

MarA is a monomeric protein composed of two helix-turn-helix (HTH) domains, with helices 3 and 6 making direct contact with the DNA (Figure 1A). Even though the N-terminal helix of MarA does not directly contact the DNA in the crystal structures, previous work, including NMR and mutagenesis studies, suggested that this structural element could be involved in DNA binding, affecting to different extents the activation of the promoters under its control ^28,30–32^. Also, the coexistence of distinct doubling resonances affecting the first ten residues of MarA indicated the co-existence of two conformations for this region of the protein ^28^.

In order to check the importance of the free movement of the N-terminal helix of MarA in the mechanism of DNA binding, we constructed two double cysteine variants to block the movement of this structural element through the formation of a disulfide bond (Figure 1). One of the variants, I13C/M47C, was designed to maintain the N-terminal helix in the same position as observed in the published MarA structures (PDB 1BL0 and 1XS9) ^27,29,36^ (Figure 1B). In the other variant, I16C/M47C, the formation of the disulfide bond immobilises the N-terminal helix outside the position observed in the PDBs (Figure 1C). This is a direct consequence of the longer distance between the two cysteines introduced at positions 16 and 47.

Wild type (WT) MarA and its variants were overexpressed, purified and tested in an electrophoretic mobility shift assay (EMSA) by using 200-bp DNA fragments harbouring the sequence of the marbox of the *marRAB* operon centred in the fragment. These experiments were performed in the presence and absence of dithiothreitol (DTT) which reduces the cysteines and breaks the disulfide bond, releasing the N-terminal helix of MarA and allowing its free movement.

The I13C/M47C variant and WT MarA were both able to bind the marbox in the absence and presence of DTT (Figure 2). By contrast, the variant I16C/M47C was only able to bind the marbox when the DTT was present, that is when the disulfide bond constraining the free movement of the N-terminal helix was broken (Figure 2). The same pattern was observed for WT, I13C/M47C and I16C/M47C when using 200-bp DNA fragments harbouring the *acrAB* marbox or 30-bp DNA fragments composed of the *marRAB* marboxes (Supplementary Figures S1 and S2 and Supplementary Table S1). The binding specificity was confirmed by using a 200-bp DNA fragment that does not include any marbox sequence as a negative control. As shown in the gels, WT MarA or its variants were not able to shift this DNA fragment (Supplementary Figure S1 and Supplementary Table S1).

**Figure 2:**
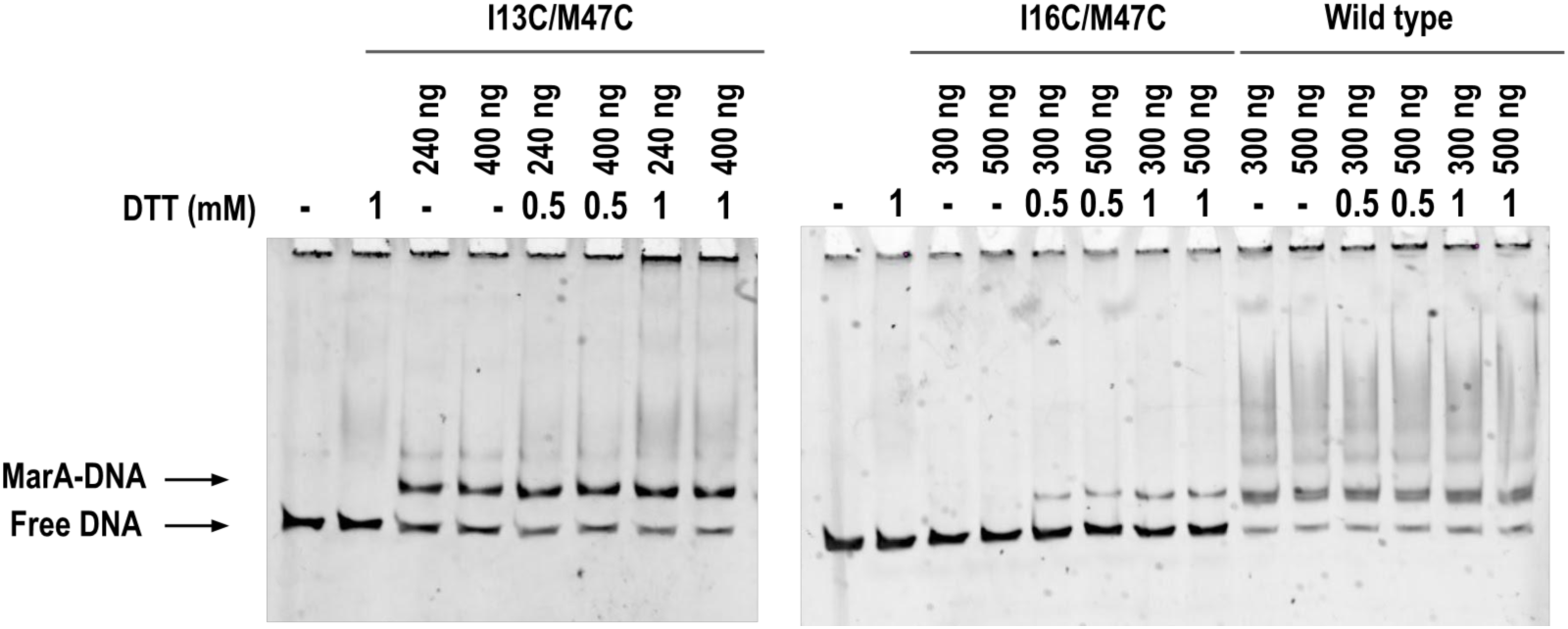
EMSA for the WT MarA and its double cysteine variants I13C/M47C and I16C/M47C in the presence and absence of DTT by using 200-bp DNA fragments containing the *marRAB* marbox. The multi-band pattern is only observed for the WT protein.

WT MarA is known to be unstable when it is not DNA-complexed and it becomes more ordered upon the formation of the MarA-DNA complex ^27,29,33^. Since the I16C/M47C variant was even more unstable that the WT (shown by the protein precipitation after overnight storage at 4°C), the EMSAs were performed with the N-terminal his-tagged proteins (Figure 2). To ensure that the His-tag did not affect the EMSA results, this tag was cleaved using thrombin and the cleaved proteins were separated by affinity chromatography. During the cleavage and subsequent separation of the His-tag and the cleaved protein, the concentration of the I16C/M47C variant dropped dramatically. Even with this, we succeed in performing the EMSA with the cleaved variants and confirmed that the pattern of bands was similar to the one observed for the His-tagged proteins (Supplementary Figure S3 and Supplementary Table S1).

Strikingly, even when the I13C/M47C variant was able to bind the marboxes in both reductive and oxidative conditions, it never showed the same multiband pattern observed for wild type MarA (Figure 2, Supplementary figures S1 and S3). To know if this pattern corresponded to non-specific binding, we performed two competitive EMSAs by using 200 bp-DNA fragments containing the *acrAB* marbox and (1) salmon sperm DNA or (2) the previously used 200-bp non-specific DNA fragment (Supplementary Figure S1) as non-specific competitors (Supplementary Figure S4A). As observed in both gels, WT MarA maintained the multiband pattern, while the I13C/M47C variant showed a single shifted band (Figure S4A and B). In this sense, the multi-band pattern exhibited by WT MarA could be related to different conformations of the WT MarA-marbox complex, or an increasing number of MarA monomers bound to the DNA, and not to non-specific binding. Further research must be done to clarify this.

To ensure that the introduction of two cysteines in MarA was not affecting the EMSAs, the S62C/L121C variant was designed as a negative control for the formation of the bonds. Specifically, in S62C/L121C, the disulfide bond blocked the movement of the C-terminal tail since the residues 62 and 121 are located in the helix 4 and the C-terminal tail, respectively. As expected, the ability to bind DNA in the presence or absence of DTT was not affected when using a 200 bp DNA fragment containing the *marRAB* sequence (Supplementary Figure S5). Finally, the formation of the disulfide bond in the N-terminal MarA variants was confirmed by mass spectrometry under oxidative and reductive conditions (see details in Supplementary data, Tables S2-S6, and Figures S6, S7).

### Intrinsic fluorescence measurements suggest that the disulfide bonds do not affect the overall fold of the N-terminal domain

To ensure that the I16C/M47C variant was not prevented from binding to the DNA by an indirect distortion of the whole N-terminal domain produced by the introduction of the disulfide bond, we performed intrinsic fluorescence measurements. WT MarA has two tryptophans in its sequence, W19 and W42, located in the N-terminal helix and helix 3, respectively. Intrinsic fluorescence is a powerful technique that allows conformational changes in proteins to be monitored by studying the tryptophan signal which is highly dependent on the local microenvironment ^37^. We reasoned that, in the absence of DTT, the maintenance of the fluorescence for W42 in the two MarA variants would indicate that the overall folding of the whole N-terminal domain is not being affected by the formation of the disulfide bond. Two peaks around 350 nm and 380 were observed in the WT MarA spectrum (Supplementary Figure S8, red line). To determine which of the two peaks corresponded to W19, we synthesised the W42F MarA variant with only one tryptophan in its sequence. As expected, this variant only showed one of the two peaks, allowing the identification of the peak corresponding to W19 (Supplementary Figure S8, orange line). When measuring the I13C/M47C and I16C/M47C variants some differences around 350 nm were observed (Supplementary Figure S8, blue and green lines). This can be explained since I16C/M47C pulls more of the N-terminal helix being W19 surrounded by different neighbour residues and generating a different microenvironment. A slight difference in the signal around 380 nm was also observed when comparing WT MarA and the variants which could be a consequence of the disulfide bond formation. In contrast, we could not see any difference in the signal of the peaks around 380 nm for the I13C/M47C and I16C/M47C variants, meaning that the microenvironment around them was similar. The fact that the two variants have the same fluorescence signal at 380 nm but show different behaviour in the EMSAs (the I13C/M47C and I16C/M47C variants able and unable to bind DNA, respectively) strongly suggests that the inability of I16C/M47C for binding the DNA after disulfide bond formation is not related to the distortion of the N-terminal domain. In this way, the inability of the I16C/M47C variant to bind DNA once the disulfide bond is formed could only be explained by the restriction of the movement of the N-terminal helix, imposed by the disulfide bond formation.

### Constraining movement of the MarA N-terminal helix reduces activation of acrAB transcription

In order to test the ability of the variants to bind the marbox and induce transcription *in vivo*, a fluorescent transcriptional reporter system was used. Specifically, it consisted of the pACYC177 plasmid containing the *E. coli acrAB* promoter followed by the gene encoding green fluorescent protein (GFP). This plasmid was named pACYC177-RS. Since we were interested in studying the activity of the variants after formation of the disulfide bond, we used shuffle T7 express *E. coli* cells that lack important reductases located in the *E. coli* cytoplasm generating an oxidative folding that allows the formation of disulfide bonds in the cytoplasm ^38^. The pACYC177 plasmids and pET-15b harbouring WT *marA*, I16C/M47C or I17C/M47C were co-transformed and *acrAB* expression was assessed by measuring GFP fluorescence.

When the cultures were grown to OD_600_ = 0.5 and induced with 0.5 mM IPTG for 90 min, 37°C, an increase in normalised fluorescence was observed for the cells containing the pET15b plasmid harbouring WT MarA in comparison with the variants or pET-15b empty vector (Figure 3A). The presence of pET-15b (empty vector) induced the synthesis of GFP slightly. The increase of fluorescence when adding empty vectors is a well-known effect that can be attributed to transcription initiation from cryptic promoter elements or regulatory elements present in the vector backbone ^39^. I13C/M47C showed an increase in the fluorescence signal showing that, even by being able to bind the DNA in the EMSAs, this variant cannot induce the transcription of the GFP protein in the reporter system as the WT MarA does. Strikingly, I16C/M47C showed a signal lower than the empty vector, suggesting that the synthesis of I16C/M47C hinders the DNA binding of the endogenous WT MarA. However, even by adding increasing concentrations of I16C/M47C to the EMSA, there was no observable decrease in the ability of the WT MarA to bind the marbox (Supplementary Figure S9). The potential inhibitory effect for I16C/M47C will be the subject of future study.

**Figure 3.**
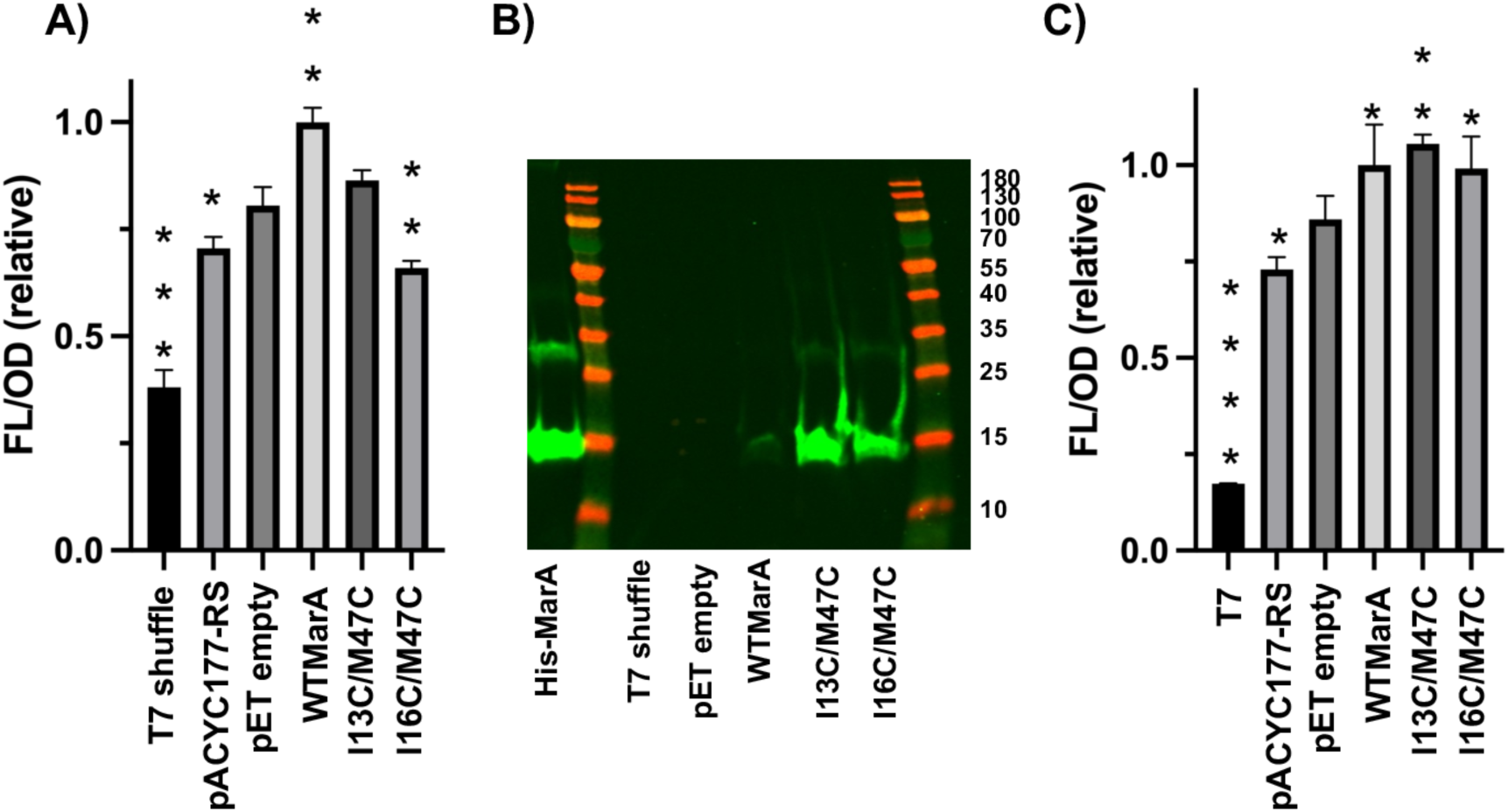
**(A)** Normalised fluorescence observed for shuffle T7 express cells, pACYC177-RS, pET-15b empty vector, WT MarA, I13C/M47C and I16C/M47C. The decrease in fluorescence exhibited by the I16C/M47C variant is statistically significant (tested by a Student’s t-test). **(B)** Western blot to detect the His-tagged MarA protein in shuffle T7 express cells, and shuffle T7 express cells harbouring the pET-15b empty vector, or pET-15b harbouring WT MarA, or I13C/M47C, or I16C/M47C. Western blotting was performed using a primary anti-His-tag mouse antibody and a secondary goat anti-mouse fluorescent antibody. MarA can be detected in the cells containing WT MarA and variants, but not in the shuffle T7 express cells or pET-15b empty vector. **(C)** Normalised fluorescence for the T7 express *E. coli* cells alone, the reporter system (pACYC177-RS) or harbouring both pACYC177-RS and pET-15b empty vector, WT *marA* gene, I13C/M47C variant, or I16C/M47C variant genes. In (A) and (C) fluorescence raw data were divided by OD and normalised such that WT MarA values were 1. The asterisks indicate statistically significant differences between pET-15b empty and the rest of the strains (tested by a Student’s t-test).

The expression of WT MarA and variants in the shuffle T7 express IPTG-induced cultures was checked by western blotting (Figure 3B). No expression of MarA was observed in shuffle T7 express cells or harbouring the pET-15b empty vector. The signal for WT MarA was much lower than for the variants. The same result was observed when using another antibody, in this case, an HRP-conjugated anti-his tag antibody, to reveal the membrane (Supplementary Figure S10). The low expression of WT MarA in shuffle T7 express cells is supported by a recent paper in which the authors observed by WB the decline of the signal corresponding to the His-tagged AraC/XylS transcriptional factor Caf1R when inducing in shuffle T7 express cells at 37°C ^40^. As the fluorescence signal for WT MarA was high in the reporter system experiment (Figure 3A), we discarded the lack of overexpression of the protein and focused on its degradation. Griffith *et al*. identified the Lon protease as the main protease involved in SoxS and MarA degradation, although they raised the possibility that a second protease also degrades MarA with very different kinetics ^26^. As explained in the main body of this paper, the shuffle T7 express *E. coli* strain is a Lon/OmpT deficient strain, so this extra protease could be acting in these cells. The WB also shows the I13C/M47C and I16C/M47C variants being less prone to degradation suggesting that the formation of the disulfide bond could avoid the degron recognition and reinforcing the idea that other non-identified proteases can degrade WT MarA by recognition of its N-terminal sequence ^26,32^.

In order to test *in vivo* if the variants can bind DNA when the disulfide bonds are not formed, we co-transformed into T7 express *E. coli* cells both, pACYC177-RS and pET15b empty or harbouring WT MarA or the I13C/M47C or I16C/M47C genes (Figure 3C). In this cell background, the reductive cytoplasm does not allow the formation of the disulfide bonds in the MarA variants. The cultures were grown to OD_600_ = 0.5 and induced with 0.5 mM IPTG for 30 min, 37°C, and an increase in normalised fluorescence was observed for the cells containing the pET15b plasmid harbouring WT MarA or its variants in comparison with pET-15b empty vector. These results show how the I16C/M47C variant can bind the *acrAB* marbox when the disulfide bond is not formed, in agreement with the results observed in the EMSAs when adding a reducing agent.

### Impact of the I13C/M47C and I16C/M47C double cysteine mutations on the dynamics and structure of free MarA

To assess the impact of the MarA I13C/M47C and I16C/M47C double cysteine substitutions on the dynamical and structural properties of MarA, we performed molecular dynamics (MD) simulations of both MarA variants in the absence of DNA, as described in the Methods section (Supplementary Figure S11). As already shown in ref ^33^, while there are some differences in mobility between free MarA and the DNA-bound crystal structure (Figure 4A and Supplementary Figure S12), the representative structure of the main cluster (containing 78% of all conformations sampled during our simulations) deviates minimally from the conformational space sampled by the DNA-bound crystal structure, so this conformation of MarA can likely easily bind the promoter sequence.

**Figure 4.**
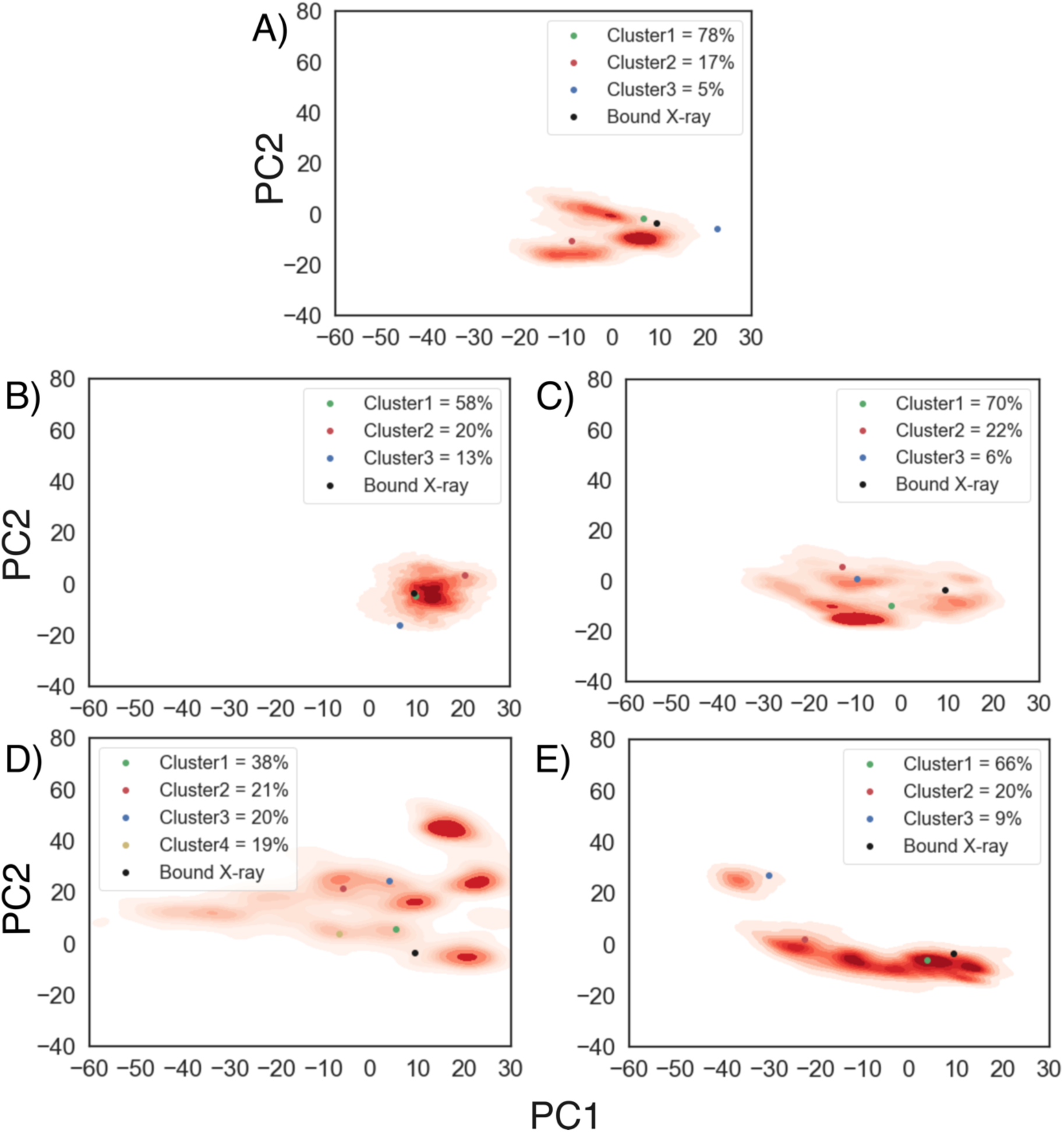
Projection of the conformational space sampled in our simulations of DNA-free wild type and MarA variants onto the first two principal components obtained from principal component analysis (PCA) on our MD simulations. Shown here are data for simulations of **(A)** wild type MarA, **(B)** the MarA I13C/M47C double variant that inserts a disulfide bridge into the system, **(C)** the MarA I13C/M47C double variant with an artificially broken disulfide bridge (named I13C/M47C^broken^) to mimic the effect of adding dithiothreitol (DTT) to the system, **(D)** the MarA I16C/M47C double variant with a disulfide bridge present in the starting structure, and **(E)** the MarA I16C/M47C^broken^. Representative structures obtained from hierarchical agglomerative clustering on the root mean square deviation (RMSD) of the Ca carbon atoms, performed as described in the Methods section (Supplementary Figure S11), are projected onto the two PCs as coloured dots. The structure of the *mar*-bound MarA crystal structure is represented by a black dot (PDB ID: 1BL0 ^27,36^).

In the case of the MarA I13C/M47C double variant (Figure 4B), PCA analysis suggests the presence of a single large basin with focused sampling of conformational space in our simulations (more so than in simulations of the wild type enzyme that lacks this disulfide bridge), with representative structures from the centroid of each cluster overlaying on that of the wild type to within a C_α_-atom RMSD of 2 Å. This suggests that the disulfide bridge already stabilizes the conformation of MarA, even in the absence of ligand, as shown in the RMSF analysis, with a reduced mobility of the N-terminal HTH motif (residues 25-51, Supplementary Figure S13). Nonetheless, when the disulfide bridge is broken, our PCA analysis shows similar mobility as observed in the wild type, although the representative structure of the main cluster slightly deviates from the conformational space sampled by the DNA-bound crystal structure (Figure 4C). In contrast, the MarA I16C/M47C variant appears to be far more conformationally diverse, both with a formed disulfide bridge (Figure 4D) and when the disulfide bridge is artificially broken in our simulations (named I16C/M47C^broken^) (Figure 4E). That is, our PCA analysis yields multiple energy basins representing different conformations of the system, irrespective of whether the disulfide bridge is formed or not. Furthermore, the broader diversity in conformations that appear to be sampled in simulations of this variant suggest a higher likelihood to sample also unproductive conformations that are unable to bind the *mar* promoter.

Following from this, visual inspection of representative structures from the centroids of each of the clusters shown in Supplementary Figure S12 shows subtle differences in structure between variants at the N-terminal HTH motif, in particular revealing structural deviations in helices H2 and H3, that partially lose their secondary structure integrity and adopt a position that would be further away from where the promoter would be expected to bind to the protein (based on the crystal structure of the DNA-bound transcription factor, see Supplementary Figure S12). This implies that for DNA binding to occur in these variants, it would require more substantial bending of the *mar* promotor for MarA to be able to establish interaction with the A-box region of *mar* promoter. Nonetheless, secondary structure analysis using DSSP (see details in Supplementary data, Figures S14 and S15) ^41,42^ shows that loss of helicity is observed in both I13C/M47C and I16C/M47C double cysteine variants, being the first fully capable of binding the *mar* sequence, and discards a change on the helicity as the reason why I16C/M47C variant is unable to bind, reinforcing the dynamical role of the N-terminal domain.

### Impact of the I13C/M47C and I16C/M47C double cysteine variants of MarA on marbox binding

To assess how the dynamic observations made in our simulations of free MarA and the I13C/M47C and I16C/M47C double cysteine variants potentially affect marbox binding, we also performed MD simulations of both MarA variants in complex with the *mar* promoter (30 bp-fragment). In all systems, MarA formed an interaction with the marbox in the starting structure of the simulations (Supplementary Figure S16A). Based on these starting structures, despite performing our sampling on the μs timescale, we did not observe unbinding of the marbox from MarA on the simulation timescales used here using cMD simulations (Supplementary Table S7), and therefore we utilized instead Gaussian-accelerated MD simulations (GaMD) to enhance the sampling of our simulations and allow for potential transitions between bound and unbound states ^33,43^.

Following from this, we observed no marbox unbinding events in our GaMD simulations of the MarA(I13C/M47C)-*mar* complex in any of the 5 replicates during 800 ns of simulation time (Supplementary Figure S17), in agreement with the experimental observation that this double variant is able to bind the marbox even in the absence of DTT. This is also consistent with the observation from our simulations of the corresponding DNA-free protein, that show that this disulfide bridge restricts the conformation of the N-terminal helix of MarA into the bound conformation observed in the crystal structure (Supplementary Figure S12). In contrast, in the GaMD simulations of the MarA(I16C/M47C)-*mar* complex, 3 out of 5 replicas lost interactions with the A-box of the *mar* promoter at the N-terminal side where the disulfide bridge is inserted, while the B-box stayed bound in all our simulations (Supplementary Figure S18). A-box unbinding observations are again in line with both the experimental data and the free MarA(I16C/M47C) simulations (Supplementary Figure S12), showing that this double cysteine variant prevents the N-terminal HTH motif of MarA to properly bend to bind the *mar* promoter. It is interesting to note here that the MarA(I16C/M47C) disulfide bridge triggered A-box unbinding but had no effect on B-box binding, while we did previously observe that introducing mutations at the nucleobases of the A-box provoked both A- and B-box unbinding on a relatively short timescale ^33^. This is interesting from a recognition mode perspective, since according to our simulations, MarA seems to not release the promoter if the sequence of A-box of the marbox is not altered, pointing to a key role of the A-box in the MarA recognition mechanism ^20^.

To further characterize the unbinding events observed in our GaMD simulations of the MarA(I16C/M47C)-*mar* complex, we used this partially unbound structure as a starting point for new cMD simulations (5 x 2.5 µs) of both the MarA(I16C/M47C)-*mar* complex, and a system where the double cysteine variant was reverted to wild type (Supplementary Figure S19). We monitored the time evolution of the insertion of both MarA HTH motifs inside the A- and B-boxes of the *mar* promoter, tracking the distances between helices 3 and 6, which are inserted inside the major groove and the base pairs at the corresponding boxes, as we did in our previous work ^33^ (*i.e.*, between Lys41-Thr52 and nucleotides 21-22 and 39-40 for the A-box and Gln91-Phe102 to nucleotides 12-13 and 49-50 for the B-box, Figure 5). On the basis of this data, we observe that the MarA(I16C/M47C) is not able to insert helix 3 inside the A-box of the *mar* promoter again, in agreement with the experimental observations, whereas the wild type reverted complex does bind the A-box of the *mar* promoter after 1.5 µs of simulation time despite starting from an unbound conformation, and this binding interaction remains stable until the end of our simulations (2.5 µs of simulation time, Figure 5).

**Figure 5.**
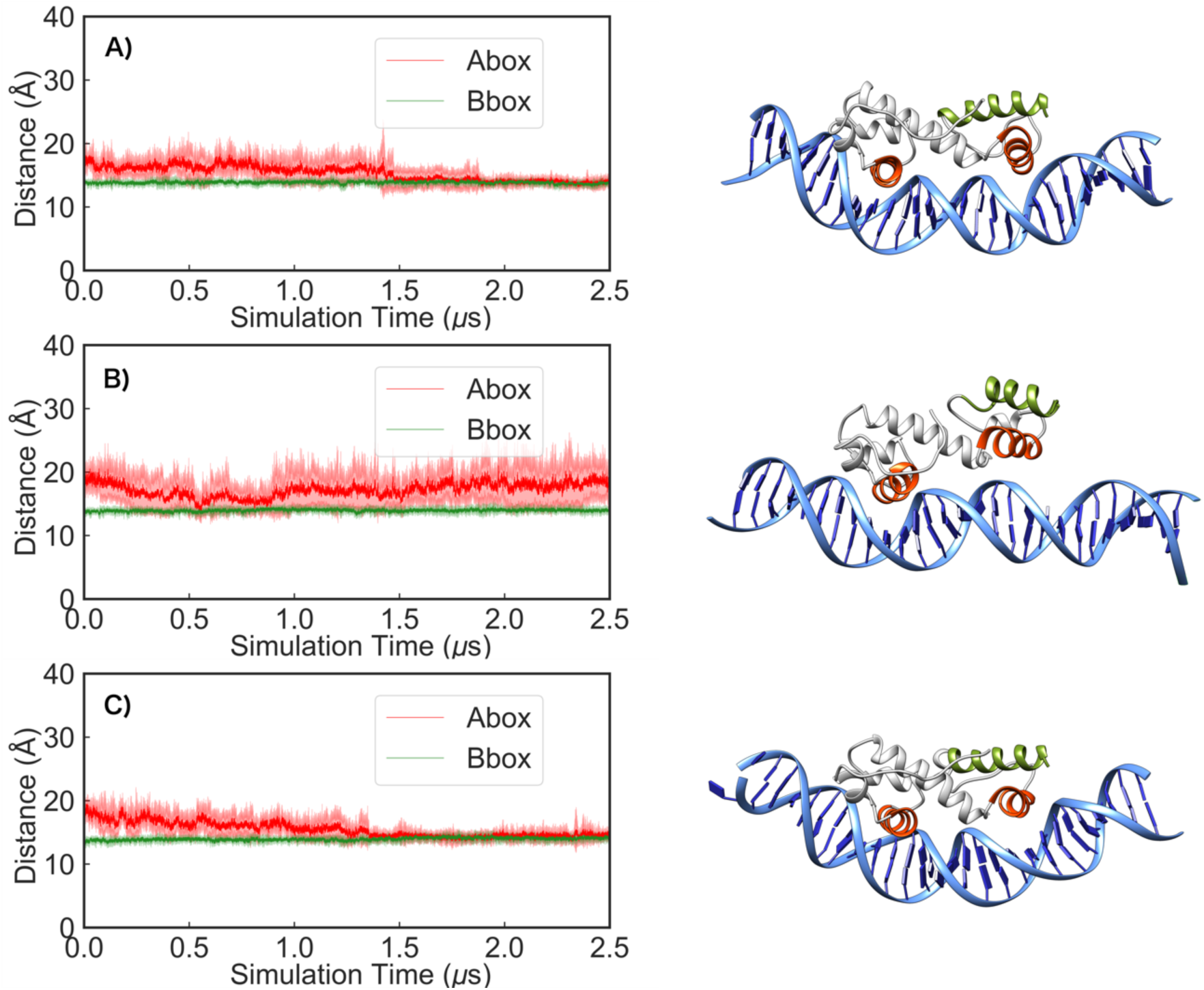
Time evolution of the distances between the helices inserted inside the major groove and the base pairs at the A- and B-boxes (shown in red and green, respectively) during 2.5 μs of conventional MD simulations of the (A) the wild type MarA-*mar* complex, and (B) the MarA (I16C/M47C) -*mar* and (C) MarA(I16C/M47C)-*mar* complexes. Note that the starting point for these simulations was the A-box unbound MarA(I16C/M47C)-*mar* complex obtained from our GaMD simulations, where in (A) the double variant was reverted to wild type and in (C) the corresponding disulfide bridge was broken by protonating each cysteine side chain. Shown here also are examples of different binding conformations during our simulations selected based on visual examination of the trajectories.

Finally, we aimed to mimic the addition of DTT to the MarA(I16C/M47C)-*mar* complex by using the same A-box unbound complex from our GaMD simulations but breaking the disulfide bridge by protonating the cysteines in both 16C and 47C positions (named I16C/M47C^broken^), and again performing (5 x 2.5 µs) of cMD simulations (Supplementary Figure S19). We tracked the same distances (Figure 5C), and observe that after 1.5 µs of simulation time, MarA is again able to interact with the A-box of the *mar* promoter, retaining a stable interaction until the end of the trajectory. Furthermore, we also performed cMD simulations (5 x 2.5 µs) using the same A-box unbound complex from our GaMD simulations, breaking the disulfide bridge and exchanging the two cysteine residues for serine to break the linkage between them while maintaining the disulfide bridge formed conformation of MarA as a starting point (Supplementary Figure S19). Again, when tracking the same distances, we observed that after around 1 µs of simulation time, MarA is again able to bind to the A-box of the *mar* promoter (Supplementary Figure S20). This strongly supports that formation of this disulfide bridge is altering the conformational dynamics of MarA, as well as the structure of the N-terminal HTH motif, in such a way that MarA is no longer able to bind the marbox of the *mar* promoter ^33,44^, and not simply due to a perturbation effect on the hydrophobic environment.

### The N-terminal helix of MarA seems to be involved in the recognition of the marboxes

MarA binds a degenerate 20-bp DNA sequence present with approximately 10,000 copies in the *E. coli* genome, most being non-functional ^21,22^. The mechanism by which MarA differentiates between the functional and non-functional marboxes remains unclear. To determine whether the N-terminal helix of MarA is involved in this mechanism of recognition, the whole N-terminal helix (ΔH1-MarA) was deleted. As the deletion of the N-terminal helix exposed the hydrophobic core of the N-terminal HTH domain, a fragment containing the Histag and ten additional amino acids was placed in its place (Supplementary Figure S21A). After purification of the overexpressed protein using the same protocol used for WT MarA, with some variations to optimise the obtaining of the DNA-free ΔH1-MarA protein (see Methods), ΔH1-MarA was found forming a complex with DNA, shown by a high 260/280 ratio (260/280 = 1.5) and by a SYBR green and Coomassie double stained gel (Supplementary Figure S21B). As expected, the pure ΔH1-MarA protein was not able to bind DNA in an EMSA (Supplementary Figure S21C).

As the same protocol for purification was used for the WT, its variants and ΔH1-MarA, these results suggest that ΔH1-MarA shows more affinity towards non-specific DNA than the WT MarA and the I13C/M47C and I16C/M47C variants. In order to test if it had less ability to differentiate the functional marboxes *in vivo*, the same reporter system as explained previously in this work was used. In this case, the IPTG-induction and overexpression of the pET-15b plasmid harbouring ΔH1-MarA did not show any increase in the fluorescent signal, indicating that ΔH1-MarA is not able to bind and induce the transcription from the *acrAB* promoter (Figure 6A).

**Figure 6.**
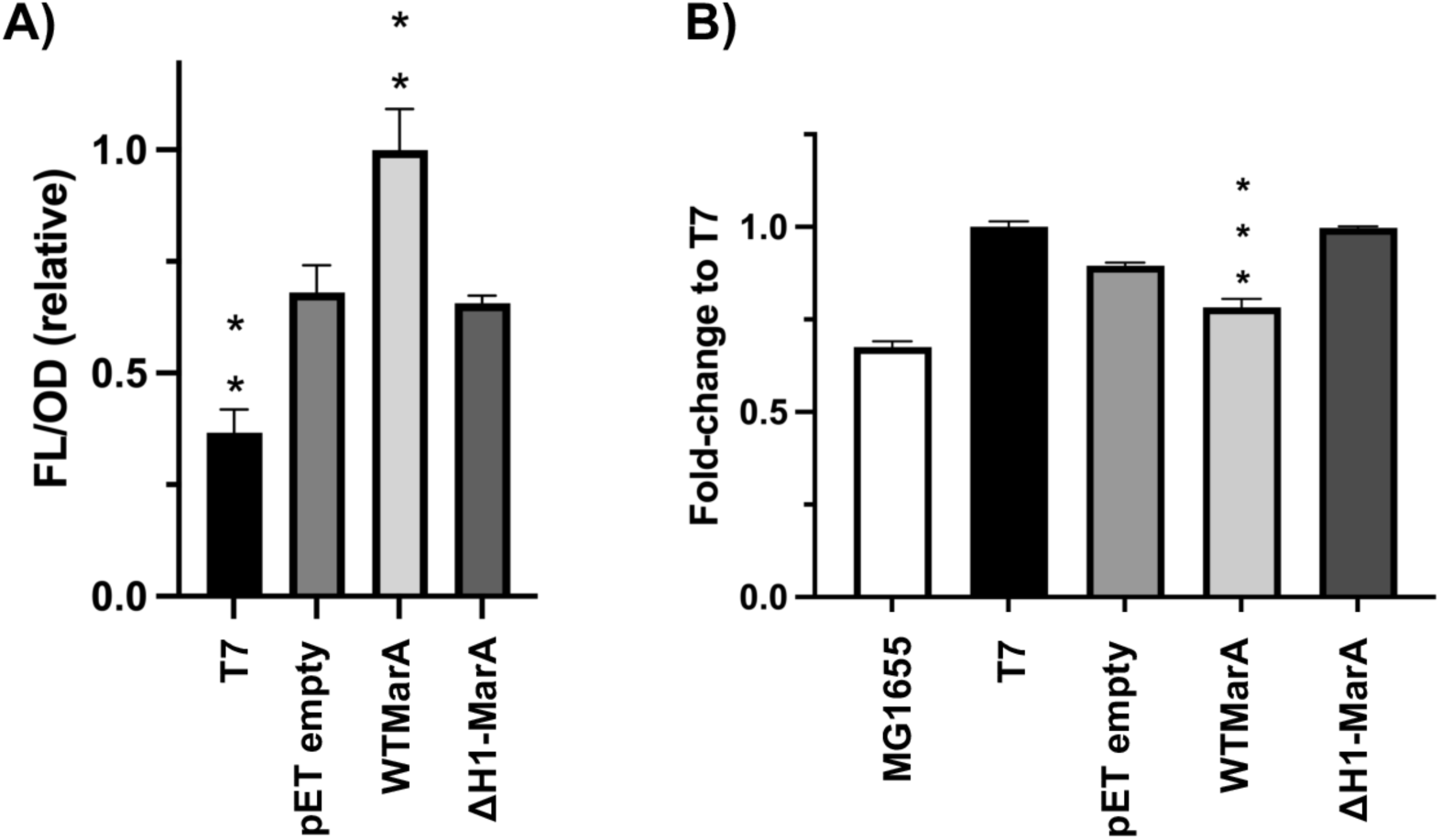
**(A)** The normalised fluorescence (FL/OD) in the reporter system shows an increase in signal after wild type (WT) MarA overexpression, but not when ΔH1-MarA is overexpressed. Fluorescence raw data were divided by OD and normalised considering WT MarA values as 1. The asterisks indicate statistically significant differences between pET empty and the rest of the strains (tested by a Student’s t-test). **(B)** Accumulation assay performed in T7 express *E. coli* cells shows that ΔH1MarA has no activity in comparison with the WT MarA, which is able to induce *acrAB-TolC* and reduce the accumulation of the dye ethidium bromide. The decrease in accumulation exhibited by WT MarA vs T7 express cells is statistically significant (shown by a Student’s t-test). MG1655 was used as a positive control.

The accumulation assay using the dye ethidium bromide showed that ΔH1-MarA was not active in the induction of *acrAB-TolC*, since it was not able to reduce the accumulation of ethidium bromide when IPTG-induced (Figure 6B). This fitted previous results, suggesting that ΔH1-MarA cannot induce the expression of the efflux pumps, maybe due to its difficulties in discerning the functional marboxes among the more than 10000 non-functional marboxes disseminated in the genome ^21,22^. Previously, it was shown that the overexpression of SoxS and MarA is toxic for the cell, with this toxicity being related to their accumulation and binding to non-functional DNA sequences located upstream of genes involved in cell growth ^26^. Further research is needed to determine if ΔH1-MarA could be toxic for the cell.

### The mechanism of inhibition involving the N-terminal helix seems to be general for the members of the AraC/XylS transcription factor family

The overlapping *marA/soxS/rob* regulon of *E. coli* contains promoters under MarA, SoxS and Rob control ^45^. These three regulators belong to the AraC/XylS family of transcription factors characterised by the presence of a segment of ∼100 amino acids that contains two HTH motifs separated by an α-helix ^46,47^. The HTH motifs are present in the DNA binding domain (DBD) and contact the DNA. The structure of these regulators showed that the DBD domain is formed by 7 α-helices, with helices 2-3 and 5-6 located in the HTH1 and HTH2, respectively, with helix 4 being a long linker, and helices 1 and 7 flanking the domain ^20,27,48^. Some of the members of the family are composed exclusively of the DBD domain (*e.g.* MarA, RamA, SoxS), but others contain extra accessory domains, called companion domain (CD), that can be placed in the N-terminus, C-terminus or in the middle of the protein (e.g. Rob or the methyl-repairing protein Ada in *E.coli* ^20,49^).

Due to the conservation of the sequence and 3D structure in AraC/XylS family members composed exclusively of the DBD (Supplementary Figure S22), we hypothesise that the mechanism of inhibition observed in MarA could also occur in other important regulatory proteins such as RamA, SoxS, or TetR. To determine if the MarA mechanism of inhibition involving the N-terminal helix could be a general inhibition mechanism for the whole AraC/XylS family including the members composed of the DBD and one or various accessory domains, new molecular dynamics simulations were performed. In these simulations, the cysteines introduced in I16C/M47C MarA were placed in the equivalent residues of Rob, and the dynamical behavior of the protein was compared to the one observed in MarA.

Rob is a 289 amino acid AraC/XylS family member composed of a N-terminal DBD and C-terminal companion domain ^20,50^. The DBD has 51% identity and 71% similarity with MarA ^47,51^. Unlike MarA and SoxS, Rob is expressed constitutively being inactive by sequestration in intracellular foci ^52^ ^53^. The crystal structures of Rob showed a different mode of DNA binding in which the DNA is unbent since only helix 3 of Rob interacts with the DNA ^20^. However, molecular dynamics simulations showed that helix 6 of Rob is conformationally dynamic, and is able to move in and out of the major groove of the DNA B-box ^33^. Recently, two cryo-EM structures have aided the understanding of the detailed molecular mechanism of Rob transcription activation ^44^.

Since the Rob crystal structure is only bound to the DNA through helix 3, and the disulfide bridge is thought to block the dynamics of the N-terminal side containing this helix, we performed two sets of simulations of Rob containing the equivalent cysteines introduced in I16C/M47C MarA, namely L10C/M41C. One set of these simulations was initiated from the Rob crystal structure (PDB ID: 1D5Y ^20,36^) with the DNA unbent and only helix 3 interacting with the DNA, and a second set was initiated with the DNA bent as in the MarA crystal structures (PDB IDs: 1BL0 and 1XS9 ^27,29,36^), allowing Rob to interact with both A- and B-boxes of the DNA (Supplementary Figure S16). We note that simulations of wild type Rob-*mar* complex in this conformation were previously performed to understand the dynamical behavior of Rob-*mar* binding recognition ^33^. Based on the data from our MarA simulations, we started directly by running GaMD on both sets of complexes. We observed no marbox unbinding events in our GaMD simulations of the Rob(L10C/M41C)-*mar* complex starting from the crystal structure binding mode in any of the 5 replicates during 800 ns of simulation time (Supplementary Figure S23). This was not surprising since when starting simulations from a complex in which the N-terminal HTH motif is singly bound to the DNA, the disulfide bridge was unlikely to alter the dynamics of Rob binding to the marbox. In contrast, in simulations of the Rob(L10C/M41C)-*mar* complex starting from initially bent DNA, 1 out of 5 replicates lost interactions with the A-box of the *mar* promoter at the N-terminal side where the disulfide bridge is inserted, while the B-box stayed bound (Supplementary Figure S24).

To further characterize the unbinding events observed in our GaMD simulations of the Rob(L10C/M41C)-*mar* complex, we used this partially unbound structure as a starting point for new cMD simulations (5 x 2.5 µs) of both the Rob(L10C/M41C)-*mar* complex, and a system where the double cysteine variant was reverted to wild type (Supplementary Figure S25). We monitored the time evolution of the insertion of both Rob HTH motifs inside the A- and B-boxes of the *mar* promoter, tracking the distances between the helices inserted inside the major groove and the base pairs at the corresponding boxes, as we did with MarA and in our previous work ^33^ (*i.e.*, between Lys35-Thr46 and nucleotides 21-22 and 39-40 for the A-box and Gln85-Phe96 to nucleotides 12-13 and 49-50 for the B-box) (Figure 7). We observe that the Rob(L10C/M41C) is not able to insert the helix 3 inside the A-box of the *mar* promoter again, as observed for MarA(I16C/M47C)-*mar*, whereas the wild type reverted complex does bind the A-box of the *mar* promoter after around 1.5 µs of simulation time despite starting from an unbound conformation, and this binding interaction remains stable until the end of our simulations (2.5 µs of simulation time, Figure 7A). Although all the simulations suggest that Rob is inhibited by the same mechanism as MarA, more experimental work is needed to confirm this mechanism of inhibition in Rob and other AraC/XylS family members.

**Figure 7.**
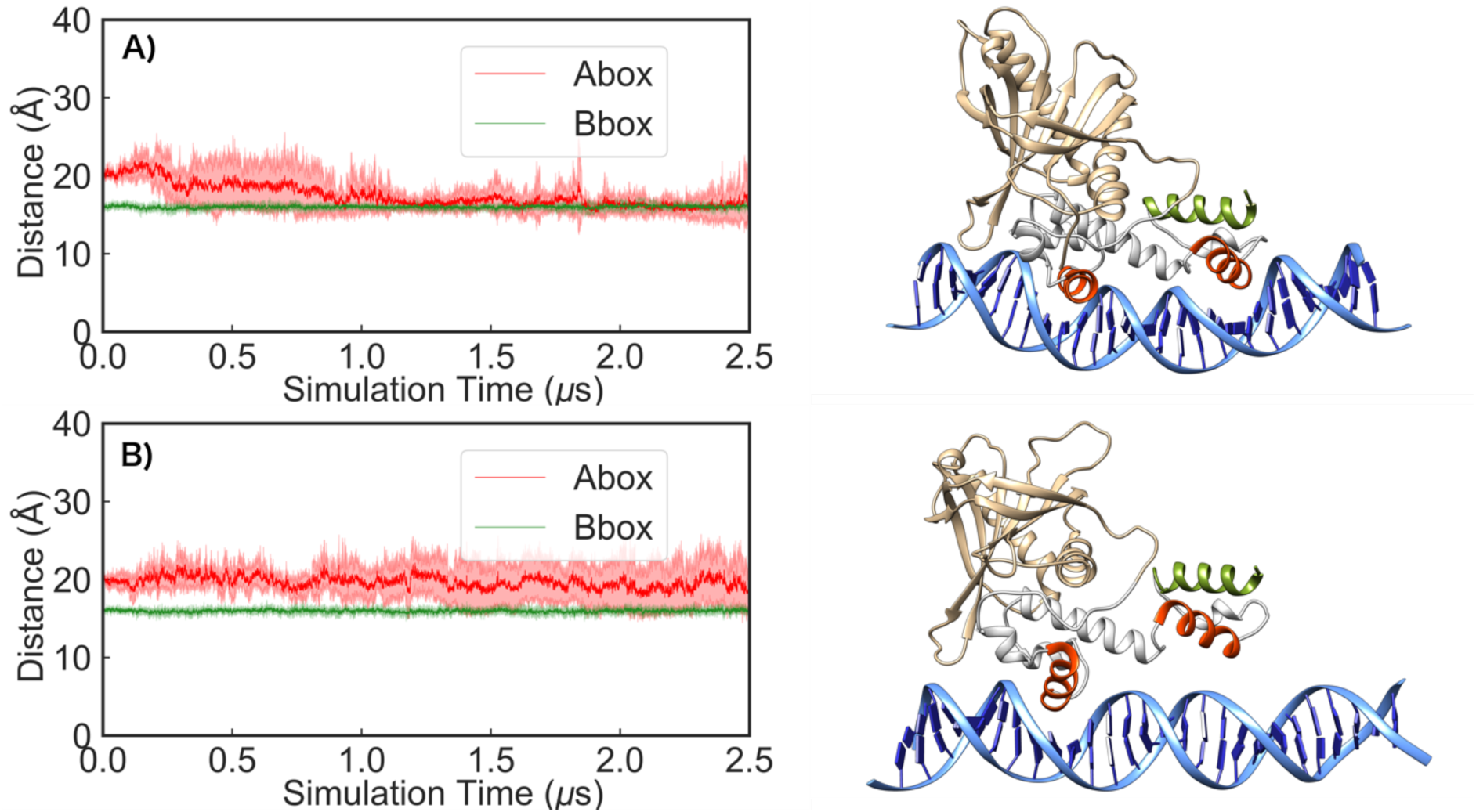
Time evolution of the distances between the helices inserted inside the major groove and the base pairs at the A- and B-boxes (shown in red and green, respectively) during 2.5 μs of conventional MD simulations of the **(A)** wild type Rob-*mar* and **(B)** Rob(L10C/M41C)-*mar* complexes. Note that the starting point for these simulations was the A-box unbound Rob(L10C/M41C)-*mar* complex obtained from our GaMD, where in (A) the double variant was reverted to wild type. Shown here also are examples of different binding conformations during our selected simulations based on visual examination of the trajectories.

## Discussion

Our study shows how the movement of the N-terminal helix of the AraC/XylS family members is key in the mechanism of DNA binding. This seems to be a general mechanism present in both, the AraC/XylS family members composed exclusively of the DBD, such as MarA, and the members having extra accessory domains, such as Rob. The identification of this new structural element, distant to the helices involved in making direct contacts with the DNA, opens the possibility for the design of new inhibitors targeting the N-terminal helix of the AraC/XylS transcription factors.

## Methods

### Bacterial strains and growth conditions

The strains used in this work were T7 express and shuffle T7 express *Escherichia coli* cells (New England Biolabs). All strains were grown in Luria-Bertani (LB) broth at 37 °C at 200 rpm. Antibiotics were purchased from Alfa Aesar (Ward Hill, MA, USA) and Merck (Darmstadt, Germany).

### Cloning and site-directed mutagenesis

The plasmid p*marA* was generated by PCR-amplification of the *marA* gene from *E. coli* MG1655 genomic DNA cloned into the *Nde*I and *HindIII* sites of pET-15b. This cloning involved the addition of a 6xHis-tag to the N-terminus of the translated protein. All site-directed mutagenesis (SDM) reactions were carried out using the Quick-Change Lightning SDM Kit (Agilent, USA), the primers listed in Supplementary Table S1, and the plasmid p*marA* as template. The double cysteine variants were prepared in two steps, by introducing the cysteines in series. The mutations were verified by sequencing (Eurofins Genomics, UK). The N-terminal deletion of MarA (called ΔH1-MarA) was obtained by amplification of the nucleotides corresponding to amino acids Glu25 to Ser129 by using the MarA_NdeI_deletionH1_F and MarA_HindIII_R primers (Supplementary Table S1). This fragment was cloned into the *Nde*I and *HindIII* sites of pET-15b as previously described for the wild type (WT) *marA*.

### Protein purification

WT MarA, its double cysteine variants I13C/M47C and I16C/M47C, and ΔH1-MarA were expressed in T7 express *E. coli* cells (New England Biolabs) grown to OD600 ≈ 0.5 and induced with 0.5 mM IPTG for 3 hours at 37 °C. The cells were harvested by centrifugation (5000 x *g*, 15 min, at 4°C), resuspended in 50 mM Tris-HCl pH 8, 1 M NaCl, 0.03 mg/ml DNase I (Sigma), supplemented with a protease inhibitor cocktail pill (Roche), and lysed by sonication. Inclusion bodies containing MarA were collected by centrifugation at 75000 x *g* for 30 minutes and washed with 50 mM Tris-HCl pH 8, 1 M NaCl, 2 M urea and centrifugated at 75000 g for 30 minutes. Inclusion bodies were solubilised with 50 mM Tris-HCl pH 8, 1 M NaCl, 7 M urea and isolated by high-speed centrifugation (75000 x *g*, 30 min). The supernatant was loaded in a 1 ml HisTrap (GE Healthcare), equilibrated with 50 mM Tris-HCl pH 8, 1 M NaCl, 7 M urea, 50 mM imidazole and eluted by increasing the concentration of imidazole up to 300 mM. Purified MarA was buffer exchanged into 50 mM Hepes pH 8, 1 M NaCl by dialysis. For the cleavage of the His-tag, thrombin sepharose beads (BioVision) were incubated with the protein overnight at room temperature. The cleaved MarA was separated by using a second round of affinity chromatography. The protein was stored at -80°C after adding 20% glycerol. As ΔH1-MarA was recovered in a complex with DNA after purification, some variations were added to the protocol such as the increase in DNase (0.06 mg/ml) and the addition of 2.5 mM MgCl_2_, 0.2 mM CaCl_2_ to improve the conditions for DNase I activity. These modifications did not allow the recovery of the pure unbound ΔH1-MarA protein.

### Electrophoretic mobility shift assay (EMSA)

For EMSA experiments, the 200-bp DNA fragments containing the *marRAB* or *acrAB* marboxes were prepared by PCR using the oligonucleotides listed in Supplementary Table S1 and *E. coli* MG1655 genomic DNA as template. The 30-bp complementary DNA fragments corresponding to the *marRAB* or *acrAB* marboxes with 5 nucleotides in each flank were purchased from Eurogentec (Seraing, Belgium, Supplementary Table S1). The DNA fragments were incubated with MarA in a buffer containing 20 mM Hepes pH 8, 100 mM MgCl_2_, 100 mM EDTA, 0.3 mg/ml BSA, and dithiothreitol (DTT) when indicated. Purified *E. coli* MarA WT or its variants were added and the mix was incubated for 20 minutes at 37°C. The samples were mixed with 6X EMSA gel-loading solution (component D, Electrophoretic Mobility Shift Assay (EMSA) Kit, Invitrogen) and loaded onto an 8% native acrylamide gel and visualised with SYBR green (Invitrogen).

### Reporter system

A reporter system built in a pACYC177 plasmid and composed of the *E. coli acrAB* promoter followed by the GFP protein (named pACYC177-RS) was co-transformed with the pET-15b empty vector, or *pmarA* harbouring the WT, ΔH1-MarA, I13C/M47C or I16C/M47C *marA* genes into T7 express or shuffle T7 express *E. coli* cells. Shuffle T7 express *E. coli* cells lack important reductases located in the *E. coli* cytoplasm generating an oxidative folding that allows the formation of disulfide bonds in the cytoplasm. We chose pACYC177 since the origin of replication is compatible with pET-15b, plasmid in which WT MarA and the variants were cloned. Shuffle T7 express cells were grown up to OD_600_ = 0.5 and induced with 0.5 mM IPTG for 90 min at 37°C while T7 express cells were grown to OD_600_ = 0.5 and induced with 0.5 mM IPTG for 30 min at 37°C. The cells were harvested by centrifugation, the pellet was resuspended with MOPS minimal medium and 190 μl of all the strains were placed in a 96 well black/clear bottom plate (Thermo Fisher) and the OD_600_ and fluorescence was measured. The fluorescence signals were normalised using the number of cells for every culture. Data presented are the mean of two biological replicates.

### Western blotting

Wild type and variants of MarA were expressed in the shuffle T7 express *E. coli* strain. Cultures were grown to an OD_600_ of 0.5 and induced with 0.5 mM IPTG for 90 min at 37°C. Cells were harvested and lysed in 50 mM phosphate buffer pH 8, 1 M NaCl, supplemented with complete EDTA-Free Protease Inhibitor tablets (Roche, Switzerland) using sonication. Membrane fractions were harvested, separated using an any kD precast SDS-PAGE gel (Biorad, UK) and transferred to a nitrocellulose membrane by using the iBlot Dry blotting system (Thermofisher Scientific, US). The membrane was rinsed in phosphate-buffered saline solution with 0.1 % Tween-20 (PBST) and blocked with PBST including 5% (w/v) milk powder for 60 min at room temperature. The membrane was then incubated overnight at 4°C with a 1:1000 concentration of anti-6×His tagged monoclonal antibody produced in mouse (ABD Serotec Biorad, UK). Then, the membrane was washed with PBST (three times for 10 min) and incubated with a 1:10000 concentration of secondary Licor IRDye 680RD-goat-anti-mouse-IgG (in TBST with 5% (w/v) milk powder for 60 min, darkness) and washed with TBS (three times for 10 min). Fluorescence detection was performed using the LiCor Odyssey (Model 9120) 3.0 imaging system.

### Ethidium bromide accumulation assay

T7 express *E. coli* cells were transformed with the pET-15b empty vector, or *pmarA* harbouring the WT or ΔH1-MarA *marA* genes. The cells were grown up to OD_600_ = 0.4 and induced by 0.5 mM IPTG for 15 min at 37°C. Later, the cells were harvested by centrifugation and the pellet was resuspended with potassium phosphate buffer pH 7, containing 1 mM MgCl_2_. After adjusting the OD_600_ to 0.2, 200 μl of each culture was inoculated into the wells of a black polystyrene microtiter tray (Greiner CELLSTAR 96 well plates flat bottom) and a 10 μl injection of 50 μg/ml ethidium bromide (EtBr) was applied to each well. The fluorescence signal was measured over 66 min at excitation and emission wavelengths of 530 and 600 nm, respectively, in a FLUOstar Optima. Data presented are the mean of three biological replicates.

### Mass spectrometry

The molecular mass of WT MarA, I13C/M47C and I16C/M47C variants, with and without a previous step of reduction with 1 mM dithiothreitol (DTT), was determined using a UHPLC-MS Nexera-i LC-2040C instrument (Shimadzu, Kyoto, Japan) with a Kinetex PS-C18 column (50 × 2.1 mm, 2.6 um; Phenomenex). The samples were eluted with a linear gradient starting at 5% of solvent B (0.08% formic acid in MeCN) and reaching 95% of solvent B over 3 min at a 0.6 mL/min flow rate, with UV detection at 220 nm. The theoretical monoisotopic masses were determined by the GPMAW software (version 12.2). The calculated and observed monoisotopic masses of each protein are shown in Supplementary Tables S2 to S6. ESI-MS spectra for WT MarA, I13C/M47C and I16C/M47C in the absence (oxidised conditions) and presence (reduced conditions) of 1 mM DTT are included in Supplementary Material.

### Intrinsic fluorescence measurements

Tryptophan fluorescence measurements were performed on a Spark multimode microplate reader (Tecan, Switzerland). The excitation wavelength was fixed at 280 nm, and the emission spectra were recorded at 25°C from 300 to 500 in a 96-well black bottom microplate (Greiner). A bandwidth of 7.5 nm was used for the excitation and emission beams, number of flashes was 30, integration time 40 µs, and gain 150. WT MarA and its variants spectra were measured at 1.2 µM in 50 mM Hepes buffer, pH 8, 1 M NaCl at room temperature. The normalised fluorescence values represent the average of three replicates that were previously corrected by subtracting the baseline corresponding to the buffer without protein.

### Bioinformatic tools

The protein sequences shown in this work were obtained by using the Uniprot Database (https://www.uniprot.org/) and the alignments were done by using Clustal Omega in the EBI (European Bioinformatics Institute) webserver (http://www.ebi.ac.uk/clustalw/) ^54,55^. The PDBs were obtained from the Protein Data Bank (https://www.rcsb.org/) ^56,57^. The figures and structural superpositions were obtained by using Chimera ^58^.

### System preparation for conventional and Gaussian accelerated molecular dynamics simulations

Starting coordinates for all the simulations in this work were taken from the crystal structure of MarA in complex with the *mar* promoter available in the Protein Data Bank (PDB ID: 1BL0 ^27,56^). Two sets of simulations were performed, one set describing free unbound MarA, and a second describing the MarA-*mar* complex, in both MarA’s wild type form, as well as the MarA(I13C/M47C) and MarA(I16C/M47C) variants. Starting structures for simulations of these variants were created by substituting the corresponding residues to cysteine using the Dunbrack 2010 Rotamer Library ^59^, as implemented in UCSF Chimera, v. 1.14 ^58^. In each case, rotamers that facilitated the formation of a disulfide bridge were selected as simulation starting points, where the disulfide bridge was imposed and thus formed at the first equilibration steps. In the case of simulations of the MarA-DNA complex, the DNA sequence in the crystal structure was slightly modified and extended at the two ends to match the 30 bp-fragment used experimentally (5’GAACCGATTTAGCAAAACGTGGCATCGGTC3’). Simulations of free MarA were initiated by manually deleting the DNA promoter from the starting structure for simulations of the analogous complexes.

Finally, in addition to the initial set of simulations starting from the DNA-bound structures of the MarA(I13C/M47C)-*mar* and MarA(I16C/M47C)-*mar* complexes (based on the wild type MarA-*mar* crystal structure), we also performed a second set of “DNA-unbound” simulations of wild type MarA (*i.e.* a C16I/C47M reversion), MarA(I16C/M47C), and MarA(I16C/M47C) with a broken disulfide bridge (named I16C/M47C^broken^), where the starting structure for each set of simulations was extracted from Gaussian accelerated molecular dynamics (GaMD) simulations of the MarA(I16C/M47C)-*mar* complex ^43^. In the latter case, we selected as our starting point a representative snapshot from the GaMD simulations where the N-terminal HTH motif of MarA has lost its interaction with the A-box of the *mar* promoter. Conventional molecular dynamics (cMD) simulations were then performed on the MarA(I16C/M47C)-*mar* complex extracted from the GaMD simulations, as well as on (1) the corresponding structure of the wild type MarA*-mar* complex, where both mutations were reverted back to wild type, (2) the corresponding complex where the disulfide bridge was broken by addition of a hydrogen atom to both C16 and C47, and (3) the corresponding complex where the two cysteine residues forming the disulfide bridge were mutated to serine (I16S/M47S). The reversion to wild type MarA was again performed using the Dunbrack 2010 Rotamer Library as implemented in UCSF Chimera ^58,59^, and the relevant rotamers selected were chosen to mimic those in the crystallographic structure of wild type MarA. Relevant data concerning the molecular dynamics simulations are available for download from Zenodo, DOI: 10.5281/zenodo.7404276.

### Conventional molecular dynamics simulations

Conventional molecular dynamics (cMD) simulations were performed following the same protocol as our prior study of MarA in complex with various DNA promoter sequences ^33^. In brief, all simulations were performed using the Amber ff14SB force field ^60^ to describe MarA and Rob, the Parmbsc1 force field to describe the DNA ^61^, and the CUDA version of the PMEMD module ^62^ of the AMBER 18 simulation package ^63^. Five independent production runs of 2.5 μs of length each with different initial velocities were performed for each system, as summarized in Supplementary Table S7. Further simulation details can be found in ref. ^33^.

### Gaussian accelerated molecular dynamics simulations

Gaussian accelerated MD (GaMD) ^43^ is an enhanced sampling method that works by adding a non-negative boost potential, that follows a Gaussian distribution, to smoothen the potential energy surface (PES) of the system, thus decreasing the energy barriers between minima and accelerating transitions between low-energy states. In this work, GaMD was applied to simulations of both the MarA(I13C/M47C)-*mar* and MarA(I16C/M47C)-*mar,* and the Rob(L10C/M41C)-*mar* complexes after an initial 700 ns of conventional molecular dynamics simulations of the DNA-bound complexes had completed. A *dual-boost* scheme was applied for the boost potential, which considers the acceleration potentials to both dihedrals and the total potential energy of the system simultaneously. The system threshold energy was set to *E* = *V*_max_ and a timestep of 4 fs was used, following from our previous conventional MD setup. The 4 fs time step was achieved *via* the hydrogen mass repartition scheme ^64^ (which involves altering the mass of hydrogen atoms to 3.024 amu to allow for the larger time step), using the PARMED module of AMBER 18 ^63^. After each cMD simulation, 84 ns of GaMD equilibration was performed in the NVT ensemble, during which time the boost potential was updated every 7.2 ns, allowing equilibrium values of the acceleration parameters *E* and *k*_0_ to be reached (for details about what these parameters mean, see ref. ^43^). Finally, 720 ns of production GaMD simulations were performed, considering 5 independent replicas for each system, comprised of 700 ns of pre-equilibration cMD followed by 84 ns of GaMD equilibration and finally 720 ns of GaMD production runs per replica. This led to a cumulative total of ∼8 μs of GaMD simulations.

### Analysis of molecular dynamics simulations

All analysis of MD simulations was performed using CPPTRAJ ^65^. Principal component analysis (PCA) was performed on the combined MD simulations of all free MarA systems by first root mean square (RMS) fitting to all C_α_ carbon atoms, using the crystal structure as a reference, and then performing PCA on all the C_α_ carbon atoms. The most-populated structures were calculated by performing agglomerative hierarchical clustering on the root mean square deviation (RMSD) of C_α_-atoms to yield 5 discrete clusters. The centroid of the most-populated cluster was then selected for further characterization. The root mean square deviations (RMSD) and fluctuations (RMSF) of the C_α_-atoms were calculated over the 5 replicas. Secondary structural propensities for all residues of free MarA were calculated using the DSSP method of Kabsch and Sander ^41^.

## Data Availability

Supplementary Data are available in a separate file. Other relevant data are available for download from Zenodo, DOI: 10.5281/zenodo.7404276

## Acknowledgements

We gratefully acknowledge all members of Dr. Blaiŕs Group (Institute of Microbiology and Infection, University of Birmingham) for valuable scientific discussions. We thank Dr. Vito Ricci and Professor Laura Piddock for sharing the pMW82 plasmid harbouring the *E. coli acrAB-GFP* reporter system and Dr Toni Todorovski for his advice in the MS section.

## Author contributions

M.C. carried out all the computational chemistry work except where noted. C. M. carried out the simulations and analysis of the free MarA systems. S.C.L.K. supervised the computational chemistry work. J.M.A.B., and E.S-V. conceived the study and designed all the lab experiments. E.S-V., and R.B-M. performed the experiments in the laboratory which were supervised by J.M.A.B., M.T. and E.S-V. Finally, M.C., R.B-M., S.C.L.K., J.M.A.B., and E.S-. V. wrote the manuscript.

