## Supplementary Data for "The N-terminal helix of MarA as a key element in the mechanism of DNA binding"

**Table of Contents**


**Supplementary Information……………………………………………………………… S3**

**Supplementary Tables ………………………………………………………………… S6**

**Supplementary Figures ……………………………………………………………… S15**

**Supplementary References …………………………………………………………… S42**

**Supplementary Information**

***Mass spectrometry shows the formation of the disulfide bond in the N-terminal MarA variants***

The formation of the disulfide bond in the N-terminal MarA variants was analysed by ESI-MS under oxidative and reductive conditions. While the wild type MarA lacks cysteine residues in its amino acid sequence, both variants contain the two introduced cysteines. It is well known that the formation of a disulfide bond results in a 2-Da reduction of molecular weight, which can be identified by Mass Spectrometry [1]. Therefore, the 2-Dalton mass variation for the I13C/M47C and I16C/M47C variants in the presence (reductive environment) and absence (oxidative environment) of 1 mM DTT would allow us to determine the oxidation state of these proteins.

The measured molecular masses for WT MarA allowed us to identify the N-terminal methionine excision, a common co-translational modification (Supplementary Figure S6) [2]. Taking this into consideration, the calculated molecular masses were in excellent agreement with the theoretical one. The average of the calculated MW was 17205.57 ± 0.87 Da and 17205.40 ± 0.70 Da in the absence and presence of 1 mM DTT respectively (the expected MW was 17205.56 Da) (Supplementary Table S2 and Supplementary Figure S7). Under oxidative conditions, the deconvoluted molecular mass from more than three peaks for I13C/M47C and I16C/M47C gave the theoretically calculated one for their respective oxidised forms (MW = 17165.44 Da) (Supplementary Table S3 and S5 and Figure S7). Additionally, under reducing conditions, we detected ions which mass correspond to the reduced form of the protein (17167.46 Da) (Supplementary Table S4 and S6 and Figure S7).

Under oxidative and reductive conditions, we could also find monoisotopic ions corresponding to the reduced and oxidised form of the protein, respectively. This result shows that disulfide bond formation/breakage does not happen at 100%, which may be due to the strength of the bond, the stability of the reducing agent or the poor accessibility of DTT due to steric impediment in the N-terminal region. The presence of monoisotopic ions corresponding to the oxidized form under reducing conditions can also be explained by disulfide bond rearrangement, as reported in studies with other proteins [1,3]. However, the predominant presence of monoisotopic ions corresponding to the respective conditions tested allowed us to confirm the existence of one intramolecular disulfide bridge in the I13C/M47C and I16C/M47C variants.

***Secondary structure analysis of free MarA (DSSP)***

Secondary structure analysis using DSSP shows that while the H3 helix (residues 41-52) maintains its helicity during simulations of wild type MarA and the I13C/M47C double cysteine variant, the helicity is relatively stable over the course of the simulations at the N-terminal portion (residues 41-47) but slightly lost at the C-terminal portion (residues 48-52) (Supplementary Figure S14). In simulations of the MarA I16C/M47C double cysteine variant, the percentage of helicity is slightly reduced, specially at the C-terminal portion (residues 45-52) although still between 60-80%. Upon breaking the disulfide bridge to mimic the effect of DTT (Supplementary Figure S14), helicity is rescued showing again same helicity as in the wild type (Supplementary Figure S14). This loss of helicity could impact the ability of MarA to bind to its promotor sequence, given the importance of R46 and W42 on the H3 helix in establishing a strong binding interaction between MarA and mar promotor [4–6], nonetheless, the fact that the loss of helicity is observed in both I13C/M47C and I16C/M47C double cysteine variants, being the first fully capable of binding the mar sequence, discards a change on the helicity as the reason why I16C/M47C variant is unable to bind and reinforces the dynamical role of the N-terminal domain.

It is worth noting that the introduction of a disulfide bridge between residues I13 and M47 does not create major structural perturbation to the MarA structure, as these residues lie in relatively close spatial proximity (Cα - Cα distance of 7Å in the wild type MarA crystal structure, PDB ID: 1BL0 [6,7]). In contrast, residues I16 and M47 are 11Å apart using the same distance metric, and therefore enforced formation of a disulfide bridge between these residues would be expected to cause greater structural perturbation. We would like to note that despite the static distance in the crystal structure between I16 and M47 alpha carbons is 11Å, the system is subjected to small fluctuations that can eventually bring the two alpha carbons into an acceptable threshold (3.0 - 7.5 Å) for a disulfide bridge formation [8] without the need for artificial displacement of the N-Terminal helix or helix 3 (Supplementary Figure S15). For comparison, we therefore also performed simulations of the DNA-free MarA I13C/M47Cbroken and I16C/M47Cbroken variants in which we manually broke the disulfide bridge (see the Methods section), in order to mimic the effect of adding DTT to the system. In the DSSP analysis (Supplementary Figure S14) we observe a rescue of the wild type helicity in the H3 helix upon removal of the disulfide bridge, while for H2, the helicity is unchanged for I13C/M47C variant and a loss of helicity is observed in I16C/M47C variant upon removal of the disulfide bridge.

**Supplementary Tables**

**Supplementary Table S1.** Primers used in this work.

| Primer name | Primer sequence (5´- 3´) |
|---|---|
| **Cloning *Escherichia coli marA* and generation of the *marA* mutants** | |
| MarA_NdeI_F | GAGGTATGCATATGTCCAGACGCAATACTG |
| MarA_HindIII_R | CAGTGACGTTGTCAAGCTTTCAACTAGCTG |
| MarA_I13C_F | CTGACGCTATTACCTGTCATAGCATTTTGGAC |
| MarA_I13C_R | CAAAATGCTATGACAGGTAATAGCGTCAGTATTG |
| MarA_I16C_F | CCATTCATAGCTGTTTGGACTGGATCGAGG |
| MarA_I16C_R | GATCCAGTCCAAACAGCTATGAATGGTAATAG |
| MarA_M47C_F | CACCTGCAACGGTGTTTTAAAAAAGAAACC |
| MarA_M47C_R | CTTTTTTAAAACACCGTTGCAGGTGCC |
| MarA_S62C_F | CAATACATCCGCTGCCGTAAGATGAC |
| MarA_S62C_R | CTTACGGCAGCGGATGTATTGGC |
| MarA_L121C_F | GAATCGCGCTTTTGTCATCCATTAAATCATTAC |
| MarA_L121C_R | GATTTAATGGATGACAAAAGCGCGATTCGC |
| W42F_F | GGTTACTCTAAATTCCACCTGCAACGG |
| W42F_R | GTTGCAGGTGGAATTTAGAGTAACCTG |
| MarA_NdeI_deletionH1_F | CTGGATCGAGGACCATATGGAATCGCCACTGTC |
| **Amplification of the DNA fragments used in the EMSAs** | |
| *acrAB*_long_prom.FOR | GCTTTTGCAATCTCGCCCAGC |
| *acrAB*_long_prom.REV | GTCCGATTTCAAATTGGTCAATGG |

| | |
|---|---|
| *marRAB*_long_prom.FOR | GTTATCCTGTGTATCTGGGTTATCAGCG |
| *marRAB*_long_prom.REV | GTTGCCCTGGCAAGTAATTAGTTGC |
| non-specific_long.FOR | GACTGACGCTCAGGTGCGAAAG |
| non-specific_long.REV | CATGCTCCACCGCTTGTGCG |
| **30-bp DNA fragments used in the EMSAs** | |
| *acrAB*_short_prom.FOR | CTTCTTGTTTGGTTTTTCGTGCCATATGTTC |
| *acrAB*_short_prom.REV | GAACATATGGCACGAAAAACCAAACAAGAAG |
| *marRAB*_short_prom.FOR | GAACCGATTTAGCAAAACGTGGCATCGGTC |
| *marRAB*_short_prom.REV | GACCGATGCCACGTTTTGCTAAATCGGTTC |

**Supplementary Table S2.** HPLC-ESI/MS for WT MarA in the absence (oxidised) and presence (reduced) of 1 mM DTT [a].

| | | WT oxidised (expected mass 17205.56 Da) | | | WT reduced (expected mass 17205.56 Da) | | |
|---|---|---|---|---|---|---|---|
| | **$[M+H]^+$ (expected)** | **$[M+H]^+$ (observed)** | **observed mass (Da)** | **Δ mass (Da)** | **$[M+H]^+$ (observed)** | **observed mass (Da)** | **Δ mass (Da)** |
| **MH9+** | 1912.73 | 1912.80 | 17206.20 | 0.64 | 1912.80 | 17206.20 | 0.64 |
| **MH10+** | 1721.56 | 1721.65 | 17206.50 | 0.94 | 1721.60 | 17206.00 | 0.44 |
| **MH11+** | 1565.15 | 1565.25 | 17206.75 | 1.19 | 1565.15 | 17205.65 | 0.09 |
| **MH12+** | 1434.80 | 1434.75 | 17205.00 | -0.56 | 1434.75 | 17205.00 | -0.56 |
| **MH13+** | 1324.51 | 1324.45 | 17204.85 | -0.71 | 1324.50 | 17205.50 | -0.06 |
| **MH14+** | 1229.97 | 1229.90 | 17204.60 | -0.96 | 1229.95 | 17205.30 | -0.26 |
| **MH15+** | 1148.04 | 1148.00 | 17205.00 | -0.56 | 1148.00 | 17205.00 | -0.56 |
| **MH17+** | 1013.09 | 1013.05 | 17204.85 | -0.71 | 1013.05 | 17204.85 | -0.71 |
| **MH19+** | 906.56 | 906.50 | 17204.50 | -1.06 | 906.45 | 17203.55 | -2.01 |
| **MH22+** | 783.08 | 783.10 | 17206.20 | 0.64 | 783.00 | 17204.00 | -1.56 |
| **MH24+** | 717.90 | 717.90 | 17205.60 | 0.04 | | | |
| **MH26+** | 662.77 | 662.80 | 17206.80 | +1.24 | 662.75 | 17205.50 | -0.06 |
| **MH30+** | 574.52 | | | | 574.55 | 17206.50 | 0.94 |

[a] The expected peaks and expected mass are similar since WT MarA lacks any cysteine in the amino acid sequence. The expected monoisotopic molecular mass values ($[M+H]^+$) were calculated by using the GPMAW software (see the material and methods section). The observed mass was calculated by using the observed ($[M+H]^+$). Δ mass indicates the variation of mass between the observed and the expected molecular mass.

**Supplementary Table S3.** HPLC-ESI/MS for I13C/M47C in the absence (oxidised) of 1 mM DTT [a].

| | I13C/M47C oxidised (expected mass 17165.44 Da) | | | | |
|---|---|---|---|---|---|
| | **$[M+H]^+$ (expected)** | **$[M+H]^+$ (observed)** | **observed mass (Da)** | **Δ mass (Da)** | **paired/not paired** |
| **$MH9^+$** | 1908.27 | 1908.20 | 17164.80 | -0.64 | -S-S- |
| **$MH10^+$** | 1717.55 | 1717.60 | 17166.0 | +0.56 | -S-S- |
| **$MH11^+$** | 1561.50 | 1561.70 | 17167.70 | +2.26 | -SH HS- |
| **$MH12^+$** | 1431.46 | 1431.90 | 17170.80 | +5.36 | N.I. |
| **$MH13^+$** | 1321.42 | 1321.60 | 17167.80 | +2.36 | -SH HS- |
| **$MH14^+$** | 1227.11 | 1227.25 | 17167.50 | +2.06 | -SH HS- |
| **$MH16^+$** | 1073.84 | 1073.65 | 17162.40 | -3.04 | N.I. |
| **$MH17^+$** | 1010.74 | 1010.70 | 17164.90 | -0.54 | -S-S- |
| **$MH18^+$** | 954.64 | 954.70 | 17166.60 | +1.16 | N.I. |
| **$MH22^+$** | 781.25 | 781.25 | 17165.50 | +0.06 | -S-S- |
| **$MH23^+$** | 747.33 | 747.35 | 17166.05 | +0.61 | -S-S- |

[a] The expected monoisotopic molecular mass values ($[M+H]^+$) were calculated by using the GPMAW software (see the material and methods section). The observed mass was calculated by using the observed ($[M+H]^+$). Δ mass indicates the variation of mass between the observed and the expected molecular mass. N.I. non-identified specie.

**Supplementary Table S4.** HPLC-ESI/MS for I13C/M47C in the presence (reduced) of 1 mM DTT [a].

| | I13C/M47C reduced (expected mass 17167.46 Da) | | | | |
|---|---|---|---|---|---|
| | **$[M+H]^+$ (expected)** | **$[M+H]^+$ (observed)** | **observed mass (Da)** | **Δ mass (Da)** | **paired/not paired** |
| **$MH9^+$** | 1908.50 | 1908.55 | 17167.95 | +0.49 | -SH HS- |
| **$MH10^+$** | 1717.75 | 1717.70 | 17167.0 | -0.46 | -SH HS- |
| **$MH11^+$** | 1561.68 | 1561.40 | 17164.40 | -3.06 | N.I. |
| **$MH12^+$** | 1431.63 | 1431.20 | 17162.40 | -5.06 | N.I. |
| **$MH13^+$** | 1321.58 | 1321.40 | 17165.20 | -2.26 | -S-S- |
| **$MH14^+$** | 1227.25 | 1227.45 | 17170.30 | +2.84 | N.I. |
| **$MH16^+$** | 1073.97 | 1073.90 | 17166.40 | -1.06 | N.I. |
| **$MH18^+$** | 954.75 | 954.70 | 17166.60 | -0.86 | -SH HS- |
| **$MH22^+$** | 781.34 | 781.35 | 17167.70 | +0.24 | -SH HS- |
| **$MH24^+$** | 716.31 | 716.45 | 17170.80 | +3.34 | N.I. |
| **$MH27^+$** | 636.83 | 636.80 | 17166.60 | -0.86 | -SH HS- |

[a] The expected monoisotopic molecular mass values ($[M+H]^+$) were calculated by using the GPMAW software (see the material and methods section). The observed mass was calculated by using the observed ($[M+H]^+$). Δ mass indicates the variation of mass between the observed and the expected molecular mass. N.I. non-identified specie.

**Supplementary Table S5.** HPLC-ESI/MS for I16C/M47C in the absence (oxidised) of 1 mM DTT [a].

| | I16C/M47C oxidised (expected mass 17165.44 Da) | | | | |
|---|---|---|---|---|---|
| | **$[M+H]^+$ (expected)** | **$[M+H]^+$ (observed)** | **observed mass (Da)** | **Δ mass (Da)** | **paired/not paired** |
| **$MH9^+$** | 1908.27 | 1908.50 | 17167.50 | +2.10 | -SH HS- |
| **$MH10^+$** | 1717.55 | 1718.0 | 17170.0 | +4.60 | N.I. |
| **$MH11^+$** | 1561.50 | 1561.90 | 17169.90 | +4.50 | N.I. |
| **$MH12^+$** | 1431.46 | 1431.50 | 17166.0 | +0.60 | -S-S- |
| **$MH13^+$** | 1321.42 | 1322.90 | 17174.70 | +19.30 | N.I. |
| **$MH14^+$** | 1227.11 | 1227.15 | 17166.10 | +0.70 | -S-S- |
| **$MH15^+$** | 1145.37 | 1145.35 | 17165.25 | -0.15 | -S-S- |
| **$MH17^+$** | 1010.74 | 1010.65 | 17164.05 | -1.35 | N.I. |
| **$MH18^+$** | 954.64 | 954.75 | 17167.50 | +2.10 | -SH HS- |
| **$MH21^+$** | 818.40 | 818.40 | 17165.40 | 0 | -S-S- |
| **$MH23^+$** | 747.33 | 747.40 | 17167.20 | +1.80 | -SH HS- |

[a] The expected monoisotopic molecular mass values ($[M+H]^+$) were calculated by using the GPMAW software (see the material and methods section). The observed mass was calculated by using the observed ($[M+H]^+$). Δ mass indicates the variation of mass between the observed and the expected molecular mass. N.I. non-identified specie.

**Supplementary Table S6.** HPLC-ESI/MS for I16C/M47C in the presence (reduced) of 1 mM DTT [a].

| | I16C/M47C reduced (expected mass 17167.46 Da) | | | | |
|---|---|---|---|---|---|
| | **$[M+H]^+$ (expected)** | **$[M+H]^+$ (observed)** | **observed mass (Da)** | **Δ mass (Da)** | **paired/not paired** |
| **$MH9^+$** | 1908.50 | 1908.50 | 17167.50 | +0.04 | -SH HS- |
| **$MH10^+$** | 1717.75 | 1717.95 | 17169.50 | +2.04 | N.I. |
| **$MH11^+$** | 1561.68 | 1562.10 | 17172.10 | +4.64 | N.I. |
| **$MH12^+$** | 1431.63 | 1431.60 | 17167.20 | -0.26 | -SH HS- |
| **$MH13^+$** | 1321.58 | 1322.90 | 17184.70 | +17.24 | N.I. |
| **$MH14^+$** | 1227.25 | 1227.20 | 17166.80 | -0.66 | -SH HS- |
| **$MH15^+$** | 1145.50 | 1145.35 | 17165.25 | -2.21 | -S-S- |
| **$MH17^+$** | 1010.86 | 1010.70 | 17164.90 | -2.56 | -S-S- |
| **$MH18^+$** | 954.75 | 954.80 | 17168.40 | +0.94 | -SH HS- |
| **$MH21^+$** | 818.50 | 818.35 | 17164.35 | -3.11 | N.I. |

[a] The expected monoisotopic molecular mass values ($[M+H]^+$) were calculated by using the GPMAW software (see the material and methods section). The observed mass was calculated by using the observed ($[M+H]^+$). Δ mass indicates the variation of mass between the observed and the expected molecular mass. N.I. non-identified specie.

**Supplementary Table S7.** Summary of all simulations performed in this work.[a]

| **Systems** | | **Individual Simulations (μs)** | **Total Simulation Time (μs)** |
|---|---|---|---|
| **Conventional Molecular Dynamics simulations** | | | |
| Free MarA | I13C/M47C disulfide bridge | 5 × 2.5 | 12.5 |
| | I16C/M47C disulfide bridge | 5 × 2.5 | 12.5 |
| | I13C/M47C | 5 × 2.5 | 12.5 |
| | I16C/M47C | 5 × 2.5 | 12.5 |
| MarA-*mar* complexes | I13C/M47C disulfide bridge | 5 × 2.5 | 12.5 |
| | I16C/M47C disulfide bridge | 5 × 2.5 | 12.5 |
| MarA-*mar* complexes from unbound structure | WT | 5 × 2.5 | 12.5 |
| | I16C/M47C disulfide bridge | 5 × 2.5 | 12.5 |
| | I16C/M47C | 5 × 2.5 | 12.5 |
| | I16S/M47S | 5 × 2.5 | 12.5 |
| Rob-*mar* complexes from unbound structure | WT | 5 × 2.5 | 12.5 |
| | I16C/M47C disulfide bridge | 5 × 2.5 | 12.5 |
| **Gaussian Accelerated Molecular dynamics simulations** | | | |
| MarA-*mar* complexes | I13C/M47C disulfide bridge | 5 × 0.804 | 4.02 |
| | I16C/M47C disulfide bridge | 5 × 0.804 | 4.02 |
| Rob-*mar* complexes | I16C/M47C disulfide bridge[b] | 5 × 0.804 | 4.02 |
| | I16C/M47C disulfide bridge[c] | 5 × 0.804 | 4.02 |
| Total simulation time | | | 166.08 |

[a] Summary of the number of individual trajectories studied per system, as well as the cumulative simulation time over all trajectories per system, and over all systems in total. All simulation times are given in μs. [b] Simulations from single B-box bound structure mimicking the crystallographic conformation (PDB ID: 1D5Y, [9]). [c] Simulations initiated from the double bound (A- and B-boxes) Rob structure.

### Supplementary Figures

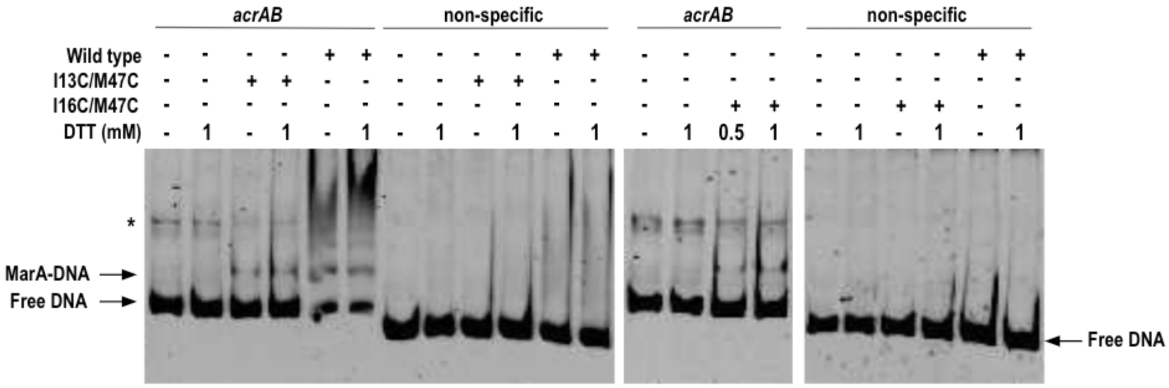


**Supplementary Figure S1.** EMSA for WT MarA and double cysteine variants I13C/M47C and I16C/M47C by using 200-bp DNA fragments harbouring the *acrAB* marbox or a non-specific sequence. On top of the lanes corresponding to the *acrAB* fragment, there is an unexpected band linked to contamination (see the asterisk).

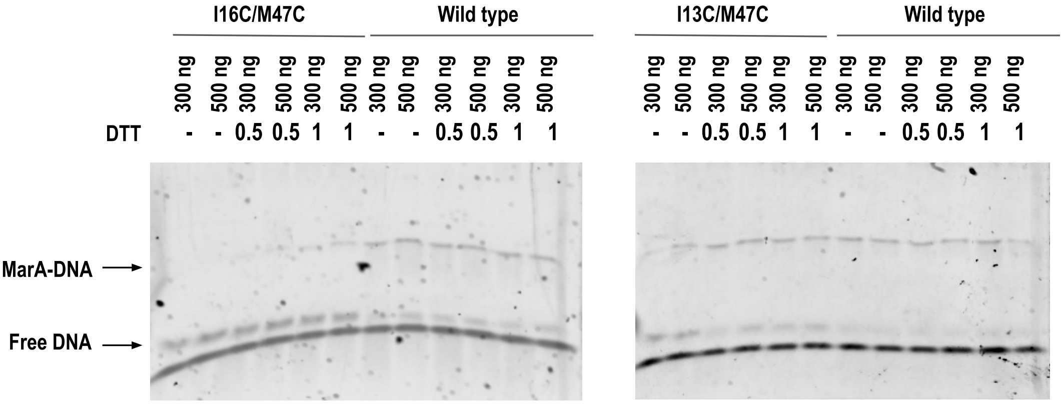


**Supplementary Figure S2.** EMSA for WT MarA and its double cysteine variants, I13C/M47C and I16C/M47C, in the presence and absence of DTT, by using 30-bp DNA fragments containing the *marRAB* marbox (see sequence in Supplementary Materials Table S1). I13C/M47C and WT MarA can bind the DNA fragments in the presence and absence of DTT. I16C/M47C only can bind DNA when DTT is present.

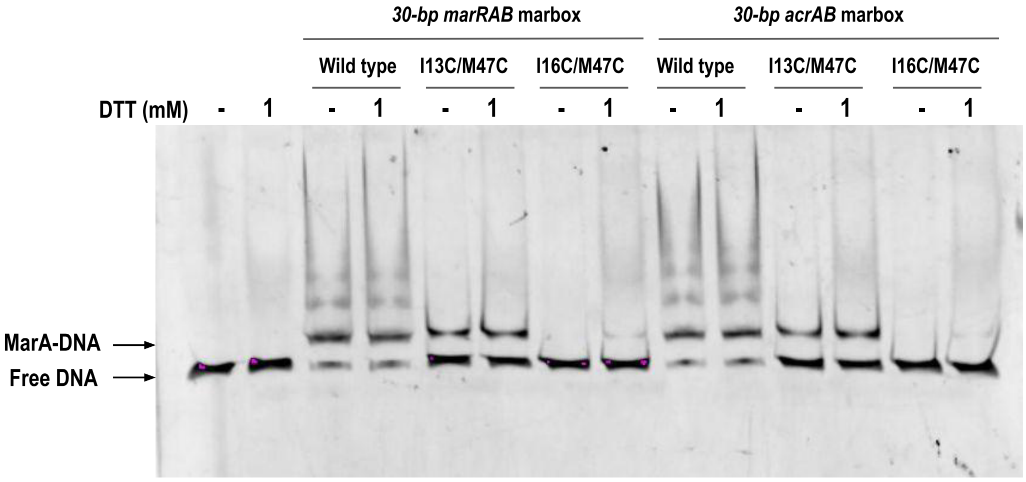


**Supplementary Figure S3.** EMSA assay by using the 30-bp *marRAB* and *acrAB* marboxes flanked by 5 amino acids in each flank (see sequences in Supplementary Materials Table S1) and the His-tag cleaved wild type MarA and double cysteine variants I13C/M47C and I16C/M47C. The pattern of bands is similar to the one obtained when working with the His-tagged proteins (Figure 2 and Supplementary Figure S1).

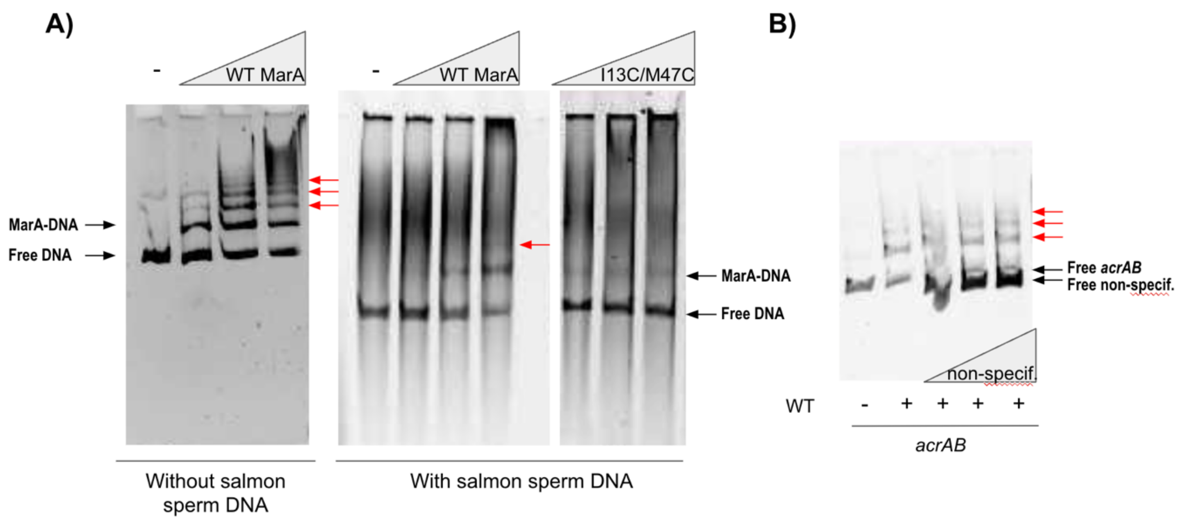


**Supplementary Figure S4.** Competitive EMSAs. **(A)** EMSA for WT MarA and I13C/M47C variant in the absence and presence of an excess of salmon sperm DNA by using 200-bp DNA fragments containing the *acrAB* marbox. As our method of detection (SYBR green) stains all the DNA present in the assay, the salmon sperm DNA was also stained complicating the visualisation of the shifted bands. **(B)** EMSA for WT MarA in the presence of an excess of 200-bp non-specific DNA. This non-specific DNA fragment was used as a competitor DNA since WT MarA and its variants were not able to shift it (Supplementary Figure S1). In both (A) and (B), the multi-band pattern exhibited by the WT remains in the presence of the non-specific competitors (red arrows). This fact indicates that the multiple bands appearing in the EMSA without salmon sperm DNA correspond to specific binding.

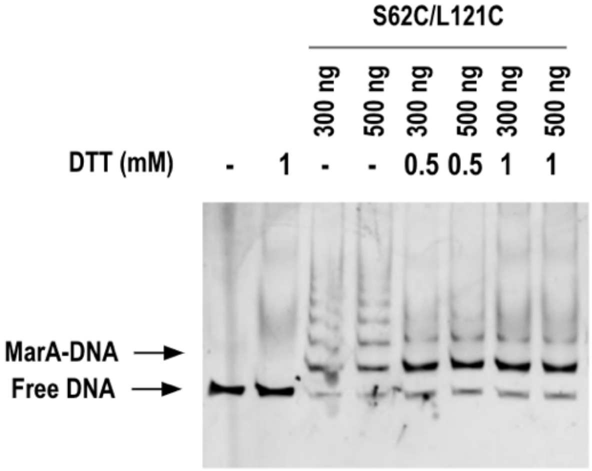


**Supplementary Figure S5.** EMSA for the double cysteine variants S62C/L121C in the presence and absence of DTT by using 200-bp DNA fragments containing the *marRAB* marbox. The disulfide bond in the S62C/L121C variant was designed to not affect the DNA binding in either the oxidised or reduced form. Specifically, in S62C/L121C, the disulfide bond blocked the movement of the C-terminal tail since the residues 62 and 121 are located in the helix 4 and the C-terminal tail, respectively. The S62C/L121C variant is able to bind the marbox in the absence and presence of DTT.

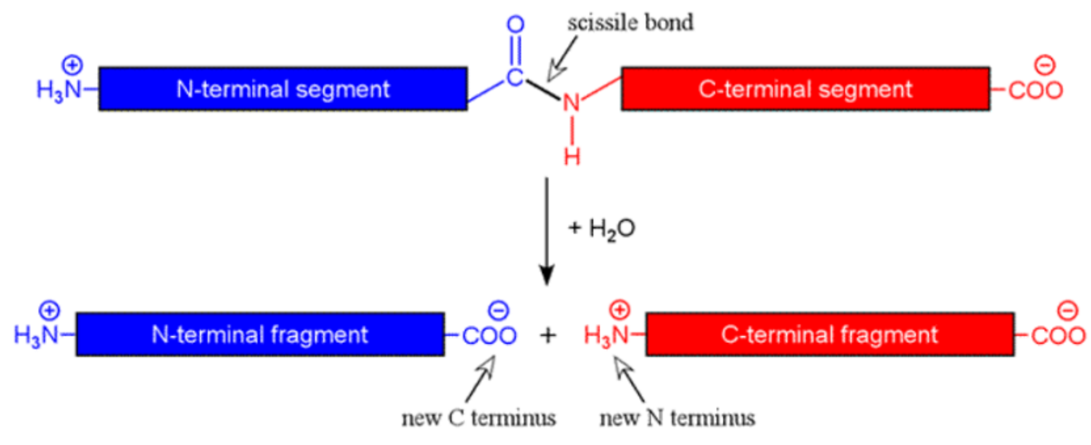


**Supplementary Figure S6.** N-terminal methionine excision, a common co-translational modification observed in WT MarA and the I13C/M47C and I16C/M47C variants. Taking the N-terminal methionine into consideration, the observed molecular masses were in excellent agreement with the theoretical one.

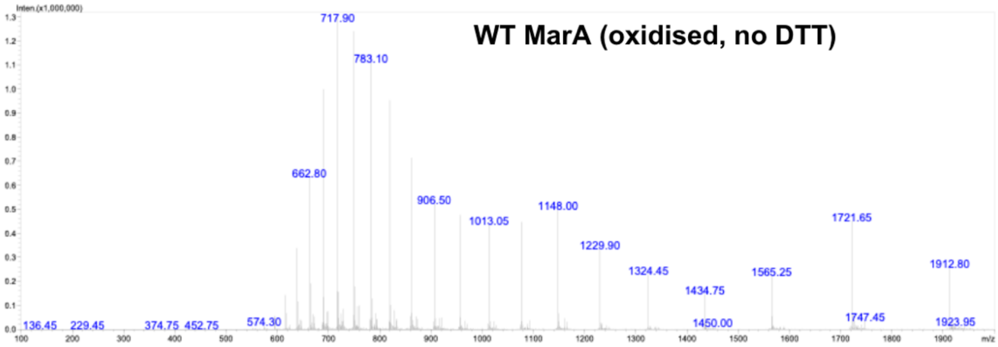
WT MarA (oxidised, no DTT)
Inten.(x1,000,000)
717.90
783.10
662.80
906.50
1013.05
1148.00
1229.90
1324.45
1434.75
1565.25
1721.65
1912.80
136.45
229.45
374.75
452.75
574.30
1450.00
1747.45
1923.95
m/z

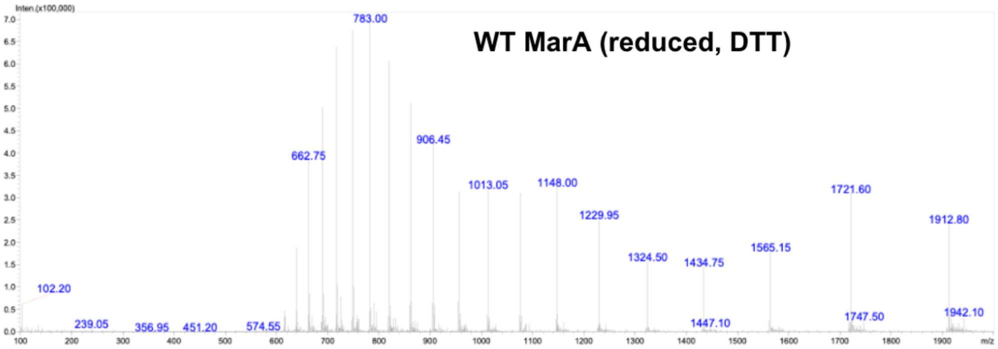
WT MarA (reduced, DTT)
Inten.(x100,000)
783.00
662.75
906.45
1013.05
1148.00
1229.95
1324.50
1434.75
1565.15
1721.60
1912.80
102.20
239.05
356.95
451.20
574.55
1447.10
1747.50
1942.10
m/z

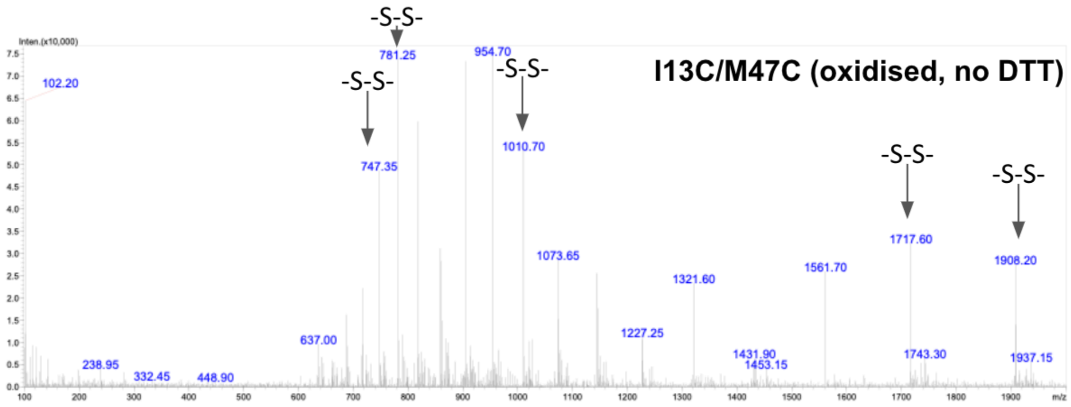
I13C/M47C (oxidised, no DTT)
-S-S-
781.25
954.70
-S-S-
747.35
-S-S-
1010.70
-S-S-
1717.60
-S-S-
1908.20
102.20
1073.65
1321.60
1561.70
637.00
1227.25
1431.90
1453.15
1743.30
1937.15
238.95
332.45
448.90
Inten.(x10,000)
m/z

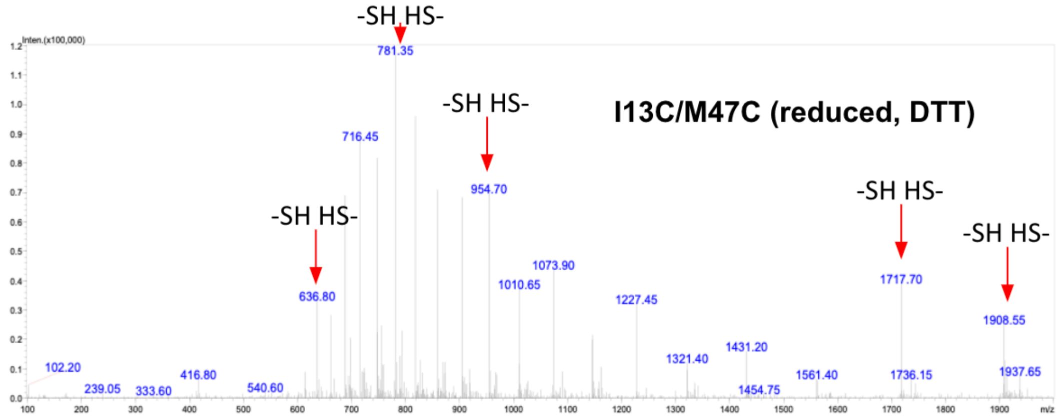
I13C/M47C (reduced, DTT)
-SH HS-
781.35
716.45
-SH HS-
954.70
-SH HS-
636.80
-SH HS-
1717.70
-SH HS-
1908.55
1073.90
1010.65
1227.45
1321.40
1431.20
1561.40
1736.15
1937.65
1454.75
102.20
239.05
333.60
416.80
540.60
Inten.(x100,000)
m/z

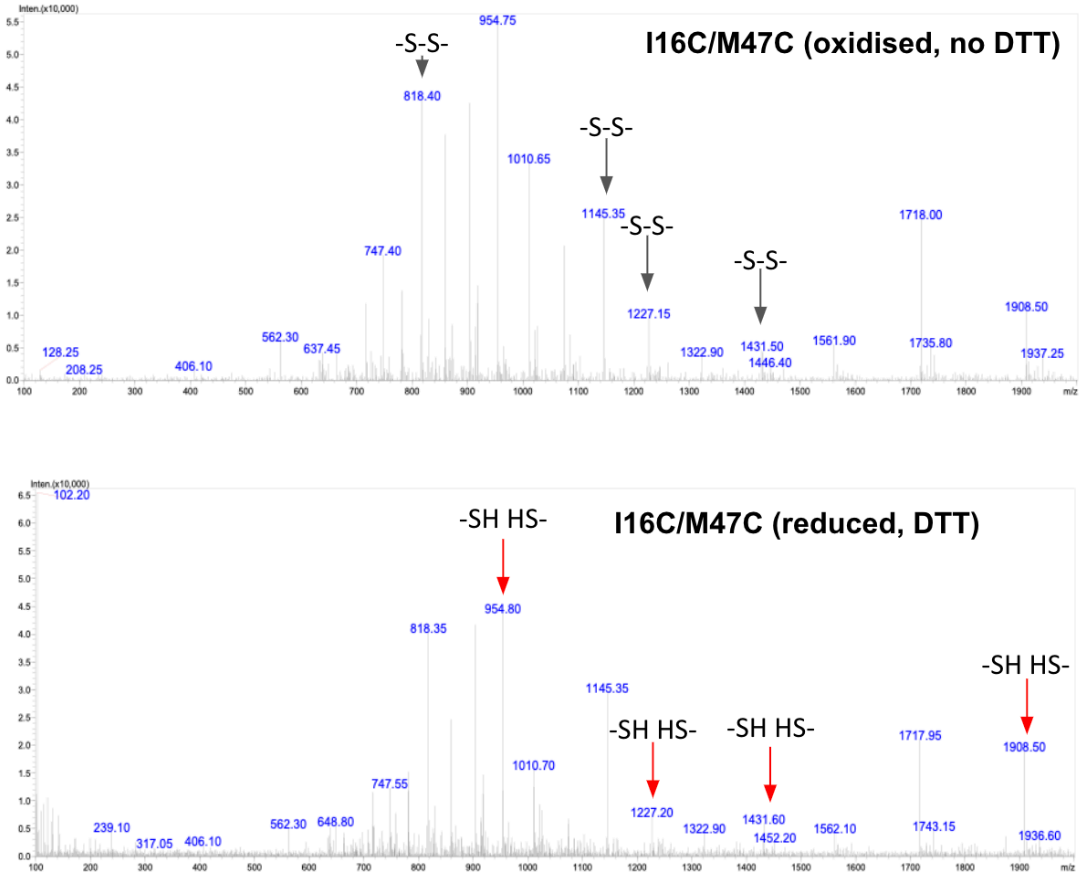


**Supplementary Figure S7.** HPLC-ESI/MS chromatograms for WT MarA, I13C/M47C and I16C/M47C in the absence (oxidised conditions) and presence (reduced conditions) of 1 mM DTT.

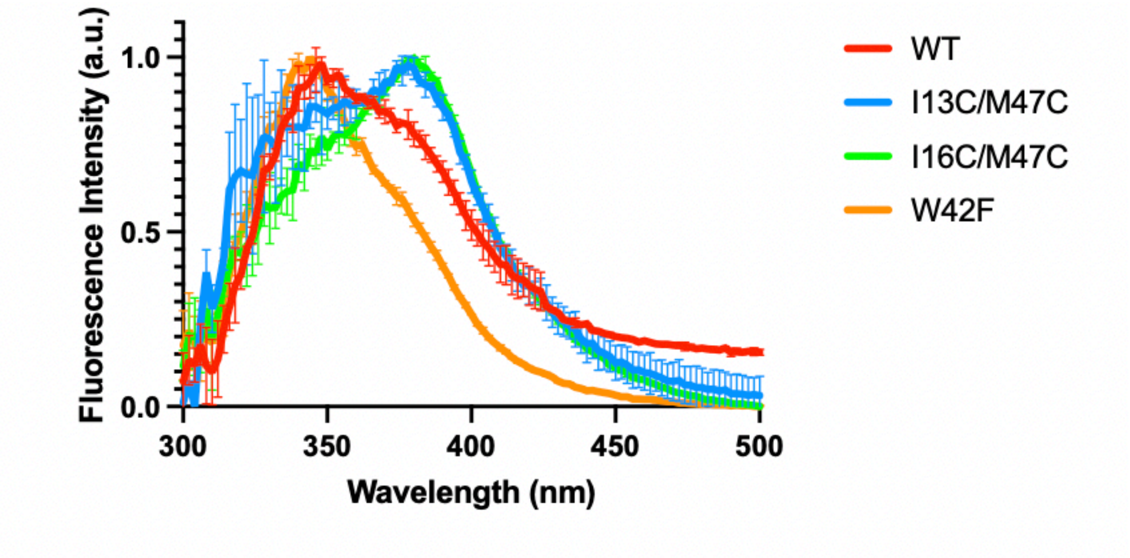


**Supplementary Figure S8.** Intrinsic fluorescence measurements of WT MarA, I13C/M47C and the I16C/M47C variants. The W42F variant was synthesised to distinguish between the signal generated by W19 (located in the N-terminal helix) and the one generated by W42 (located in the helix 3, which contact directly the DNA (Figure 1)). The I13C/M47C and I16C/M47C variants showed differences in the peak related to W19 (around 350 nm) but not related to W42 (around 380 nm). This suggests that the inability of I16C/M47C to bind DNA after disulfide bond formation does not relay on the distortion of the overall fold of the N-terminal domain, but in the immobilisation of the N-terminal helix.

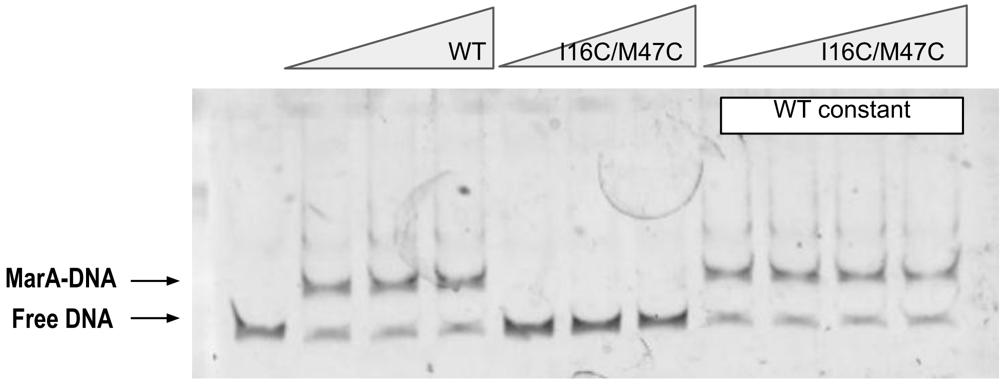


**Supplementary Figure S9**. EMSA using 200 bp-DNA fragments containing the *marRAB* marbox. Wild type (WT) MarA was tested alone (range 200 – 600 ng), I16C/M47C alone (range 100 – 300 ng) and both variants together (increasing I16C/M47C concentrations (100 – 350 ng) maintaining the WT concentration constant (200 ng)). In this assay, DTT was not added. The addition of increasing concentrations of I16C/M47C does not seem to affect WT MarA binding to the marbox.

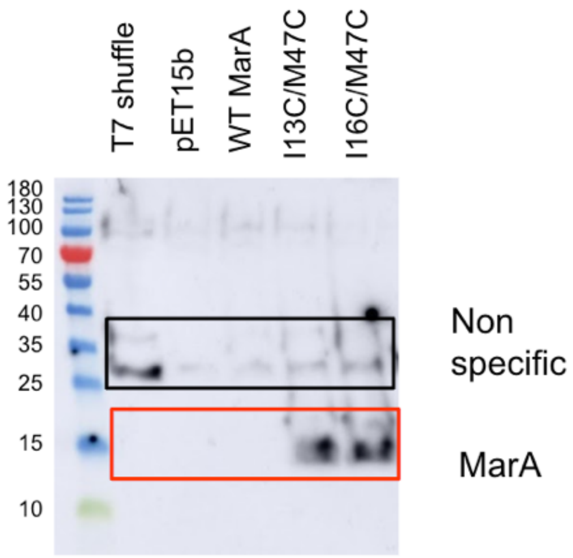


**Supplementary Figure S10.** Western blot to detect the His-tagged MarA protein in shuffle T7 express cells, and shuffle T7 express cells harbouring the pET-15b empty vector, or pET-15b harbouring WT MarA, or I13C/M47C, or I16C/M47C. Western blotting was performed using an HRP-conjugated anti-His tag antibody. MarA can be detected in the cells containing the variants, but not in the shuffle T7 express cells, pET-15b empty vector or WT MarA. The non-specific bands appearing in all the tested strains indicates that this antibody was less selective than the one used in Figure 3B.

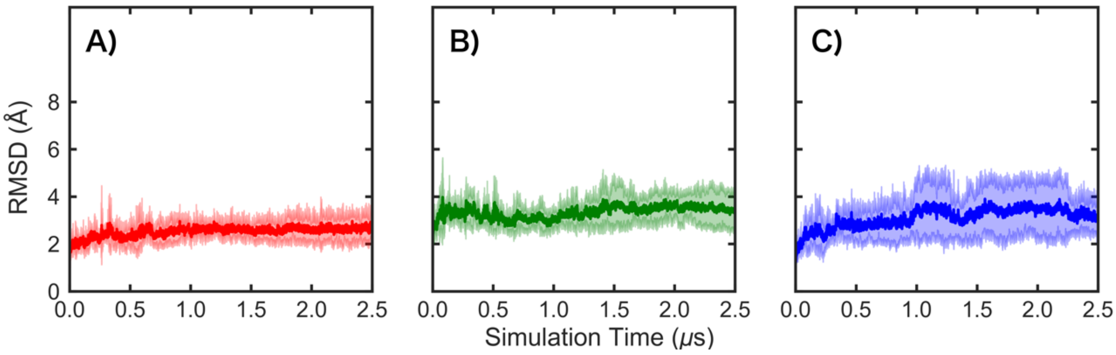


**Supplementary Figure S11.** Root mean square deviation (RMSD, Å) of all protein backbone atoms of **(A)** wild type free MarA, and the **(B)** I13C/M47C and **(C)** I16C/M47C variants. The RMSDs were calculated over 5 independent 2.5 μs molecular dynamics simulations, relative to the initial constructs. The solid lines denote the average RMSD over all replicas, and the shaded lines show the standard deviations over the different individual replicas.

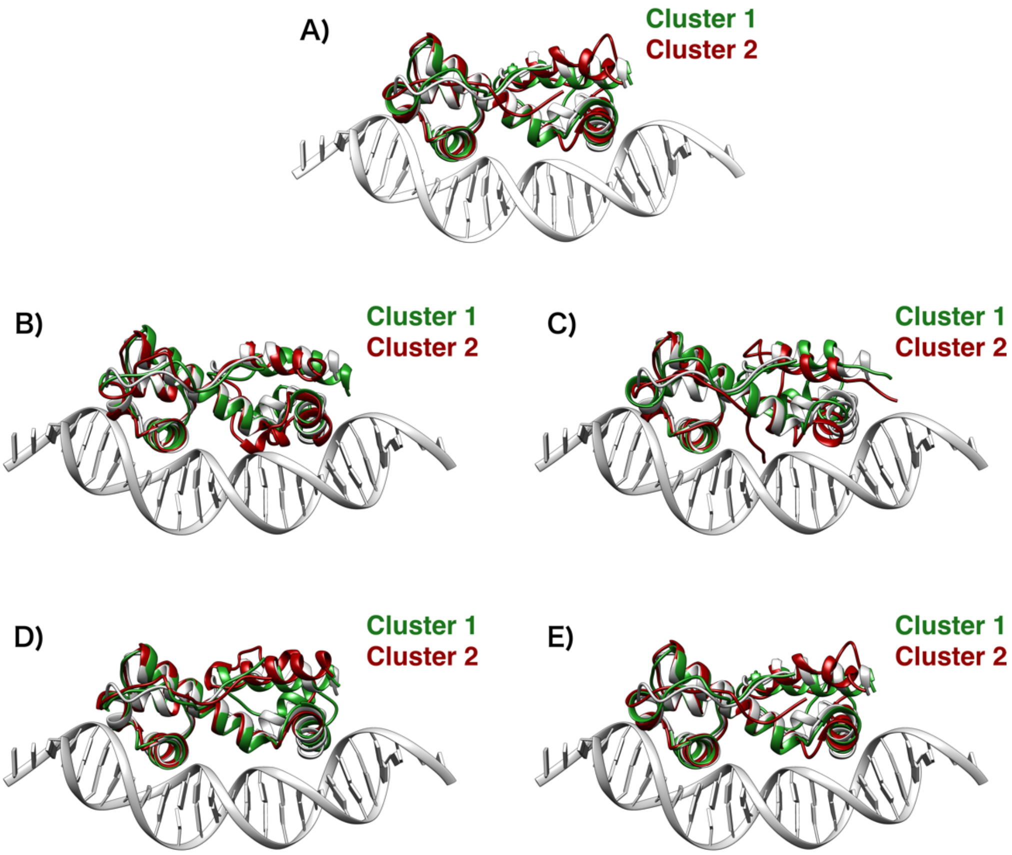


**Supplementary Figure S12.** Centroids of the first two clusters obtained from hierarchical agglomerative clustering on simulations of free MarA, with root mean square deviation (RMSD) clustering performed over 5 independent 2.5 μs molecular dynamics simulations of **(A)** wild type, **(B)** I13C/M47C, **(C)** I16C/M47C and **(D, E)** I13C/M47C and I16C/M47C MarA without the disulfide bridge present, respectively. An overlay of MarA crystal structure bound to the DNA (PDB ID: 1BL0, [7]) is shown in white. The clusters were extracted as indicated in the Methods section.

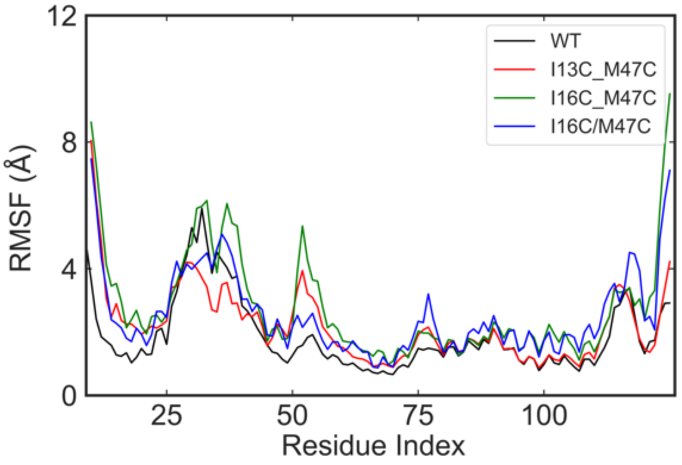


**Supplementary Figure S13.** Root mean square fluctuations (RMSF, Å) of the Cα-atoms of DNA-free MarA variants considered in this work, calculated over five independent 2.5 μs MD simulations (Supplementary Materials Table S2).

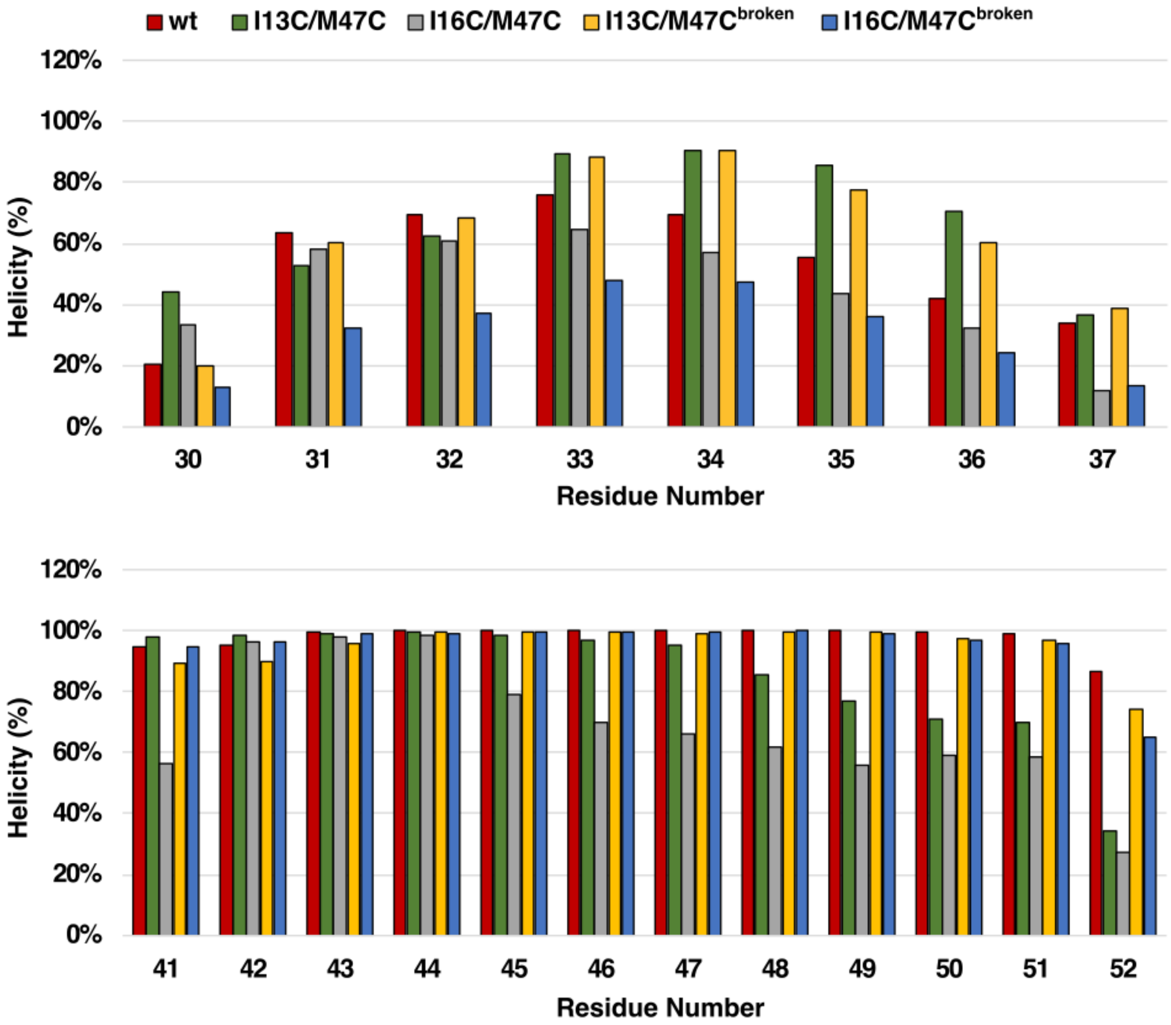


**Supplementary Figure S14.** Helicity propensities for all residues belonging to the helix-turn-helix (HTH) motif encompassing helices H2 and H3 of free wild type MarA (red), the I13C/M47C double variant MarA with a disulfide bridge (green), the I16C/M47C double variant MarA with a disulfide bridge (grey) and the I13C/M47C and I16C/M47C double variants MarA with an artificially broken disulfide bridge to mimic the effect of adding DDT to the system (yellow and blue, respectively).

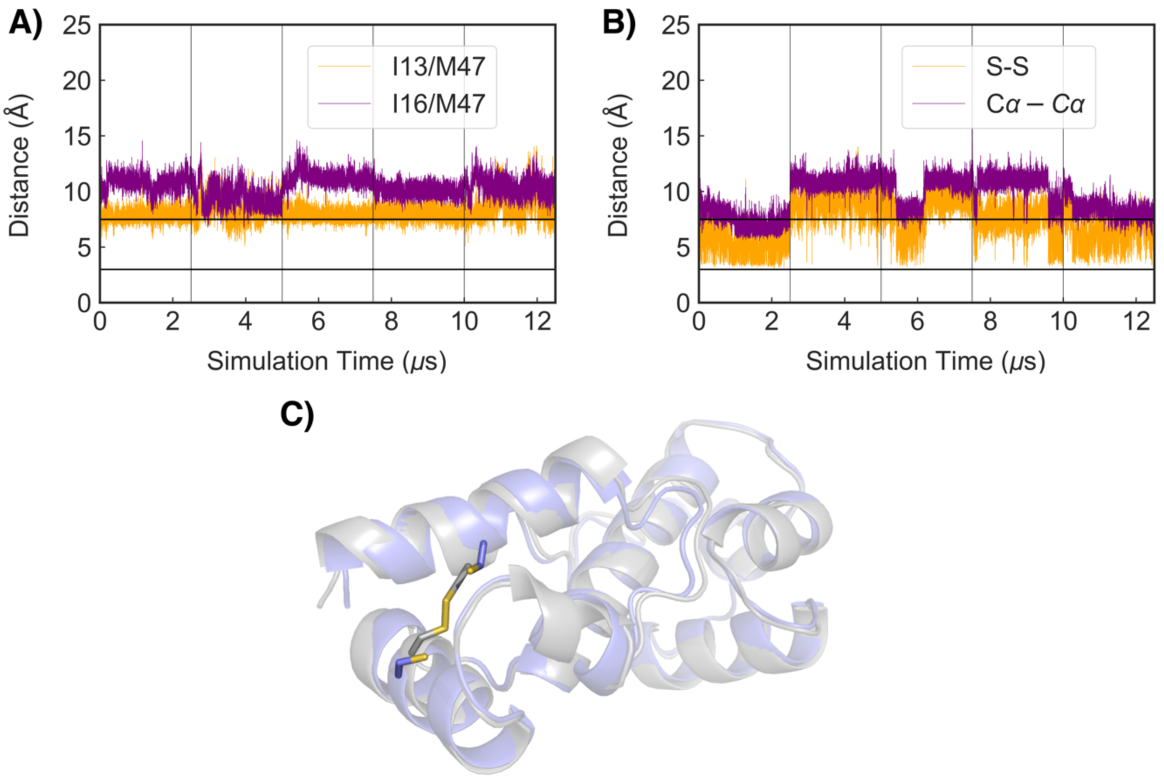


**Supplementary Figure S15.** Time evolution of the distances between the $C_\alpha$ carbon atoms of **(A)** I16/M47 and I13/M47 in wild type MarA, and **(B)** time evolution of the distances between alpha carbons and sulphur atoms of I16C/M47C, where the disulfide bridge is not present because the cysteines are reduced to SH. **(C)** Overlay of the structures of I16C/M47C MarA before and after the disulfide bridge formation.

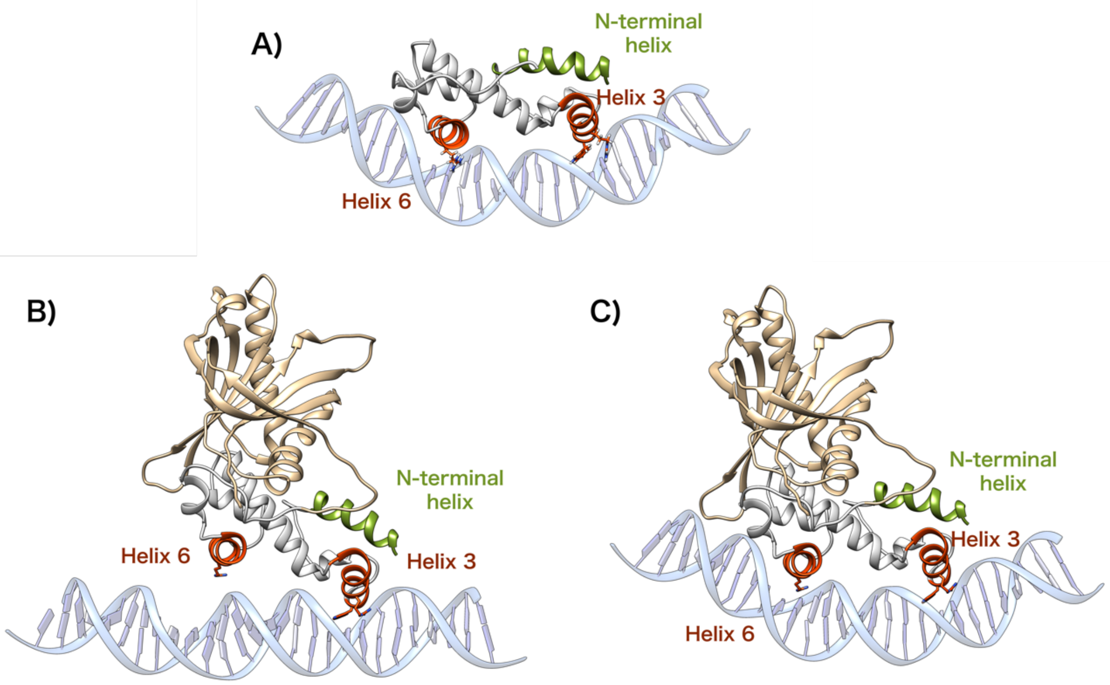


**Supplementary Figure S16.** Starting structures used in our molecular dynamic simulations of wild type and variant complexes of **(A)** MarA-*mar*, **(B)** Rob- *mar* and **(C)** Rob- *mar* with bent DNA. Helices 3 and 6 are depicted in orange-red while the N-terminal helix, where the disulfide bridge between helix 3 is created, is shown in green. Residues displaying key interactions with the DNA nucleobases are shown as sticks.

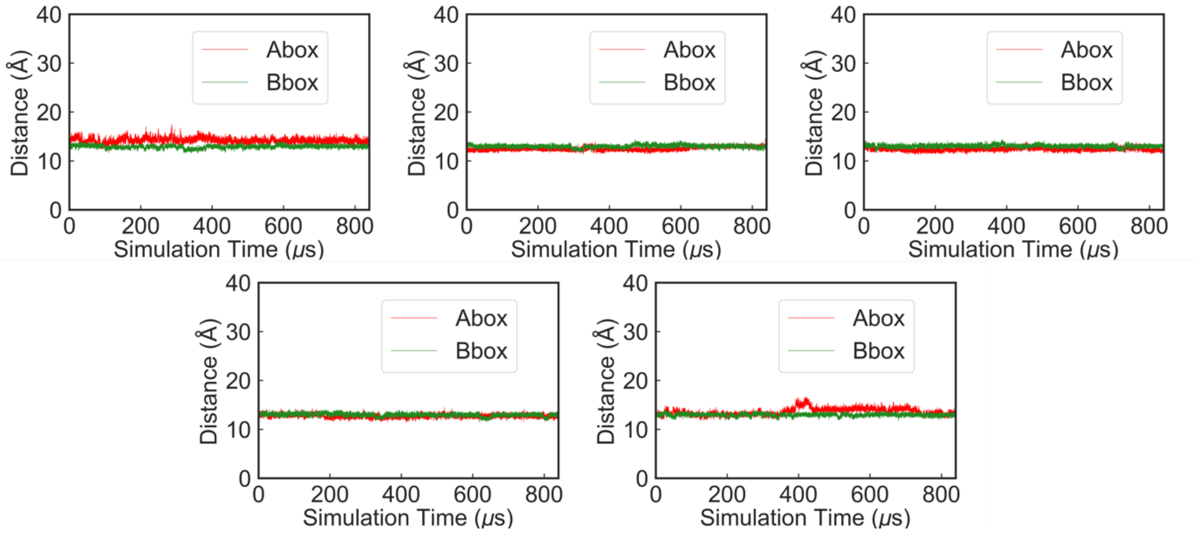


**Supplementary Figure S17.** Time evolution of the distances between helices 3 and 6 of MarA(I13C/M47C), which are inserted inside the major groove of mar, and the base pairs at the A- and B-boxes, respectively, during 5 individual replicas of GaMD simulations of the MarA(I13C/M47C)-*mar* complex.

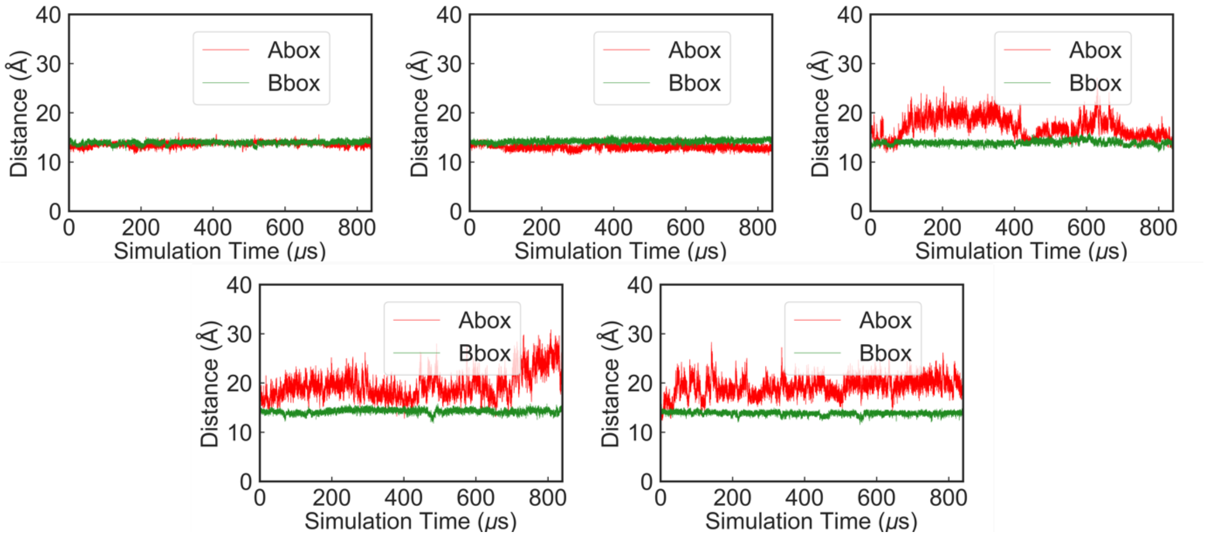


**Supplementary Figure S18.** Time evolution of the distances between the helices 3 and 6 of MarA(I16C/M47C) which are inserted inside the major groove of mar, and the base pairs at the A- and B-boxes, respectively, during 5 individual replicas of GaMD simulations of the MarA(I16C/M47C)-*mar* complex.

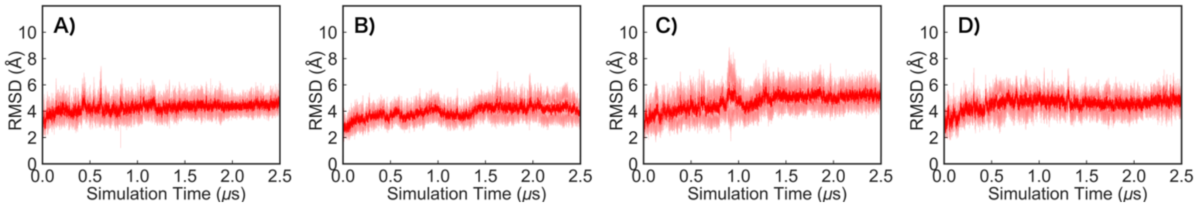


**Supplementary Figure S19.** Root mean square deviation (RMSD, Å) of all protein backbone atoms of MarA in complex with mar promoter. Shown here are data from simulations of the **(A)** wild type, the **(B)** I16C/M47C disulfide bridge variant, the **(C)** I16C/M47C variant without the disulfide bridge and **(D)** the I16S/M47S double variant, calculated over 5 independent 2.5 μs molecular dynamics simulations, and relative to the initial constructs. The solid lines denote the average RMSD over all replicas, and the shaded lines show the standard deviations over the different individual replicas.

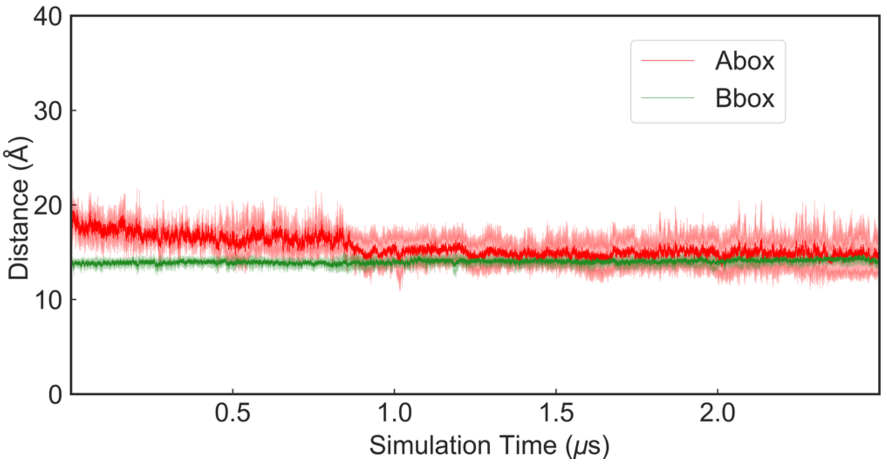


**Supplementary Figure S20.** Time evolution of the distances between the helices inserted inside the major groove and the base pairs at the A- and B-boxes during 2.5 µs of conventional MD simulations of the MarA(I16S/M47S)-*mar* complex. Note that the starting point for these simulations was the A-box unbound MarA(I16C/M47C)-*mar* complex obtained from our GaMD simulations, where the corresponding disulfide bridge was broken and each cysteine was exchanged by a serine.

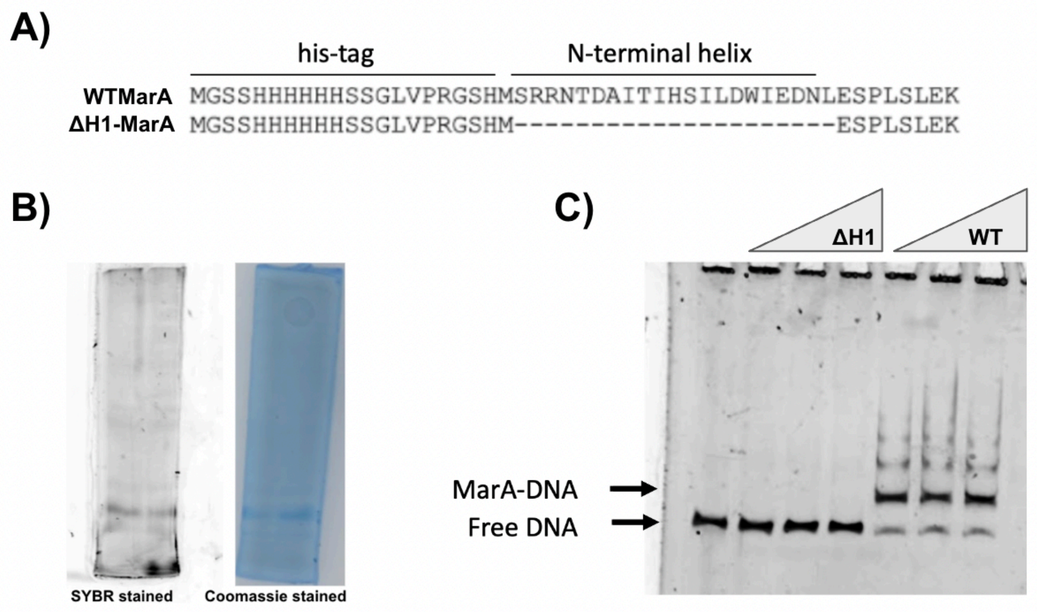


**Supplementary Figure S21.** ΔH1-MarA. **(A)** Alignment of the His-tagged wild type (WT) MarA N-terminal region and ΔH1-MarA. The deleted amino acids and the His-tag that occupies their position can be observed. **(B)** Purified ΔH1-MarA in an acrylamide gel stained with SYBR green (DNA detection) and Coomassie (protein detection). Both DNA and protein are in the same band, suggesting their co-existence as a MarA-DNA complex. **(C)** EMSA with the WT MarA protein and the ΔH1-MarA. As expected, ΔH1-MarA was not able to bind the 200-bp DNA fragment containing the *marRAB* marbox.

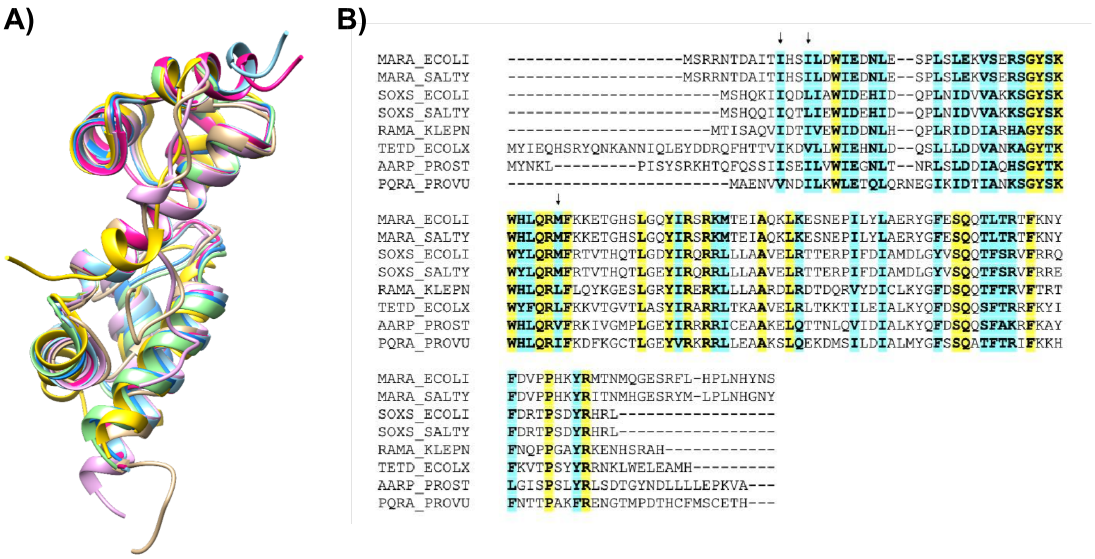


**Supplementary Figure S22.** Comparison with other AraC/XylS family members consisting exclusively of the DNA binding domain (DBD). **(A)** Superposition of MarA, SoxS, RamA, TetD, Aarp, PqrA (structures predicted by AlphaFold with a high level of confidence (pLDDTs between 83 and 93) [10,11]). **(B)** Alignment of the amino acid sequences showing the conservation of the key residues in this work. The conserved and similar residues are shadowed in yellow and cyan, respectively.

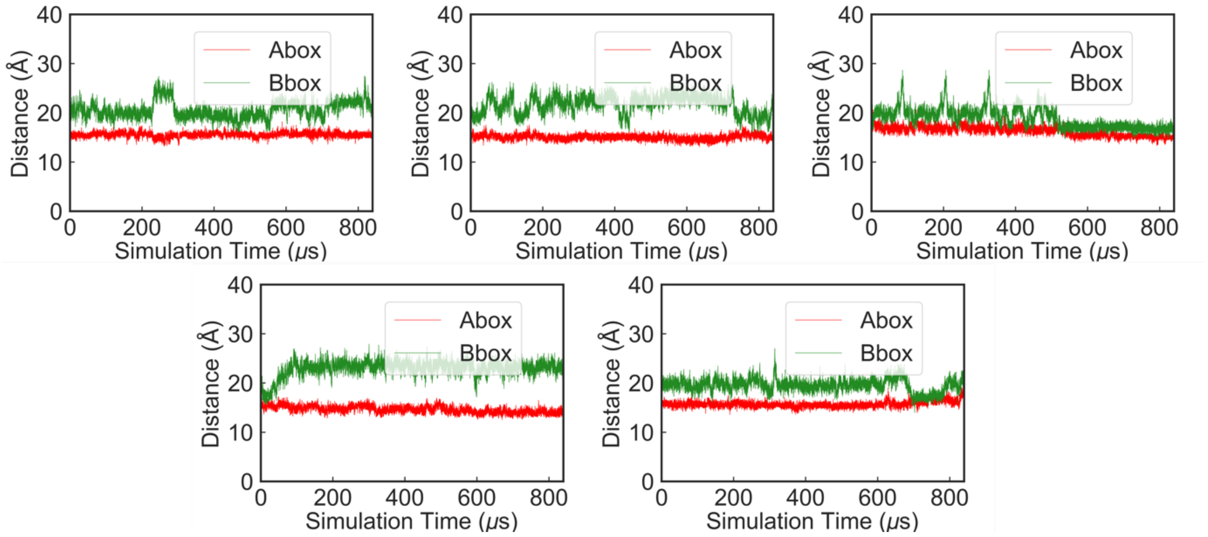


**Supplementary Figure S23.** Time evolution of the distances between the helices 3 and 6 of the Rob(L10C/M41C) variant, which are inserted inside the major groove of *mar*, and the base pairs at the A- and B-boxes, respectively, during 5 individual replicas of GaMD simulations of the Rob(L10C/M41C)-*mar* complex. Simulations were initiated from the crystallographic DNA binding mode (see the main text for details).

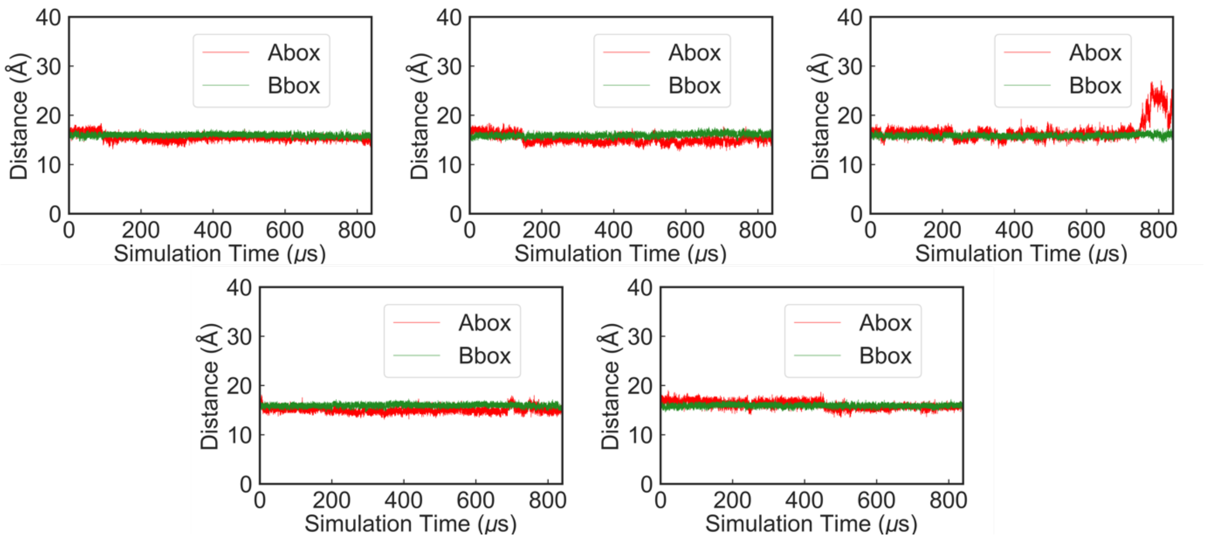


**Supplementary Figure S24.** Time evolution of the distances between helices 3 and 6 of the Rob(L10C/M41C) variant, which are inserted inside the major groove of *mar*, and the base pairs at the A- and B-boxes, respectively, during 5 individual replicas of GaMD simulations of the Rob(L10C/M41C)-*mar* complex. Simulations were initiated from the double bound DNA binding mode, with the DNA bent as in the MarA crystal structure (see the main text for details).

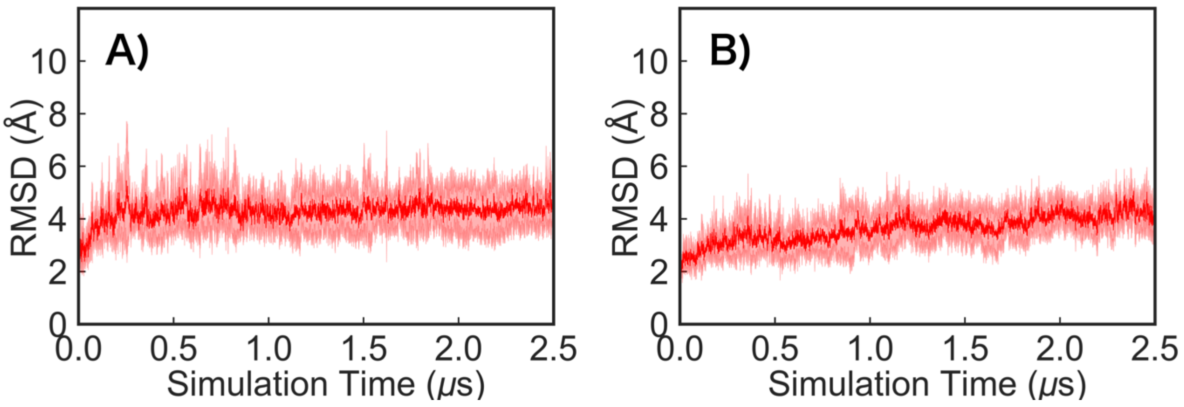


**Supplementary Figure S25.** Root mean square deviation (RMSD, Å) of all protein backbone atoms of Rob in complex with the *mar* promoter. Shown here are data from simulations of **(A)** wild type Rob and **(B)** the L10C/M41C disulfide bridge variant, calculated over 5 independent 2.5 μs molecular dynamics simulation. RMSD are calculated relative to the initial constructs. The solid lines denote the average RMSD over all replicas, and the shaded lines show the standard deviations over the different individual replicas.
